# Pre-Analytical Drivers of Bias in Bead-Enriched Plasma Proteomics

**DOI:** 10.1101/2025.05.07.652495

**Authors:** Kathrin Korff, Johannes B. Mueller-Reif, Dorothea Fichtl, Vincent Albrecht, Alicia-Sophie Schebesta, Ericka C.M. Itang, Sebastian Virreira Winter, Lesca M. Holdt, Daniel Teupser, Matthias Mann, Philipp E. Geyer

## Abstract

Bead-based enrichment has emerged as a promising strategy to improve depth in plasma proteomics by overcoming the dynamic range barrier. However, its robustness against pre-analytical variation has not been sufficiently characterized. Here, we systematically evaluate five plasma proteomics workflows—including three bead-based methods and two conventional protocols—using controlled spike-ins of low-abundance proteins and defined cellular contaminants. We find that bead-based approaches enhance detection of low-abundance proteins but can be highly susceptible to systematic bias from platelet and PBMC contamination, even at low levels. This can easily inflate results by thousands of proteins, likely accounting for some of the very high literature-reported numbers. In contrast, a perchloric acid-based workflow shows notable resistance to erythrocyte and platelet-derived contamination. We further investigate how centrifugation conditions, anticoagulant choice, and buffer-bead combinations modulate contamination profiles and demonstrate that bias can partially be mitigated by optimized sample handling. In total, we identify more than 13,000 different protein groups from various conditions, including cellular components from the circulating proteome. Our results provide a quantitative framework for assessing workflow performance under variable sample quality and offer guidance for both biomarker discovery and quality control in clinical proteomics studies.

## Introduction

Blood plasma is one of the most valuable and widely collected biofluids for clinical diagnostics, with protein-based laboratory tests constituting the largest proportion of routine clinical assessments (Anderson & Anderson, 2002; Geyer *et al*, 2017). The plasma proteome is of fundamental importance for monitoring health status, pathogenic processes and treatment (FDA-NIH Biomarker Working Group, 2016; Ignjatovic *et al*, 2019). Classical examples of protein biomarkers are cardiac troponins for myocardial infarction or liver enzymes like ASAT and ALAT for hepatic dysfunction, representing well-established diagnostic tests. Most of the currently used biomarkers were introduced decades ago and there remains a significant need for the discovery of novel biomarkers for many diseases and treatments (Anderson *et al*, 2013; Deutsch *et al*, 2021).

The complexity of the plasma proteome poses tremendous analytical challenges for biomarker discovery. Only twenty proteins account for approximately 99% of the total protein content, creating an extraordinary dynamic range that spans more than twelve orders of magnitude (Anderson & Anderson, 2002; Lee *et al*, 2011). This dynamic range presents a fundamental challenge for comprehensive proteome analysis, particularly in detecting low-abundance proteins such as tissue leakage markers or signal molecules, which are important for disease understanding and have great potential as novel biomarkers (Li *et al*, 2024).

Mass spectrometry (MS)-based proteomics is a rapidly advancing technology that increasingly provides unbiased and hypothesis-free analysis of complex samples (Aebersold & Mann, 2016; Guo *et al*, 2025). Recent technological advances including data-independent acquisition (DIA) alongside fast and high-resolution mass analyzers have significantly boosted the achievable plasma proteome depth (Lancaster *et al*, 2024; Serrano *et al*, 2024).

As importantly, emerging sample processing techniques have expanded the range of quantifiable proteins by directly addressing the dynamic range issue. This includes bead-based techniques enriching a subset of the plasma proteome in a bead-corona (Blume *et al*, 2020; Wu *et al*, 2024) or precipitation-based approaches such as perchloric acid precipitation (Viode *et al*, 2023, 2024; Albrecht *et al*, 2025). These methods aim to reduce the dominance of high-abundance proteins while being sufficiently robust to maintain quantitative integrity of proteome-wide analyses, thereby enabling the exploration of low-abundance species.

While these technological improvements have enhanced plasma proteome coverage, successful biomarker discovery requires careful consideration of many factors beyond the number of identified plasma proteins. The proteomics community has established guidelines for biomarker development that address quality standards and emphasize proper cohort selection to ensure statistical significance and clinical relevance (Hoofnagle *et al*, 2016; Mischak *et al*, 2010; Skates *et al*, 2013; Surinova *et al*, 2011). However, there remains a critical gap in systematically assessing proteome-wide effects of pre-analytical sample handling. Considering that plasma samples are often collected during routine clinical practice with variable processing conditions, these pre-analytical factors are known to significantly impact study outcomes (Rai *et al*, 2005; Schrohl *et al*, 2008; Timms *et al*, 2007). While this variability affects single samples and patients it is particularly problematic in case-control studies, where systematic differences in sample collection or processing between groups can lead to false biomarker candidates. Of particular concern are contaminations from blood cells, especially platelets and erythrocytes, which can introduce systematic bias in clinical studies. We previously reported that approximately half of the published plasma proteomics studies may be affected by such sample-related quality issues (Geyer *et al*, 2019).

In this study, we present a comprehensive evaluation of five distinct plasma proteomics workflows with regards to pre-analytic factors: a neat plasma workflow for simple and rapid analysis, a precipitation-based approach using perchloric acid with neutralization (PCA-N) (Albrecht *et al*, 2025), and three distinct bead-based enrichment methods using magnetic or non-magnetic beads with representative surface chemistries. Through carefully designed spike-in experiments using various blood cell types as well as the entire yeast proteome, we systematically assess the ability of each workflow to detect low-abundance proteins. We also characterize their susceptibility to common sample contaminants, including platelets, erythrocytes, and peripheral blood mononuclear cells (PBMCs). Additionally, we investigate how standard clinical centrifugation protocols and blood collection tubes affect plasma proteome analysis, revealing critical insights into the relationship between sample preparation parameters and proteome composition. We further ask if compromised samples can be ‘rescued’ by additional processing steps. Our findings shed light on the potential and limitations of bead-based workflows and offer practical guidance for both prospective study design and retrospective quality evaluation of archived samples.

## Results

### Study overview and investigated plasma proteomics workflows

We designed a comprehensive experimental framework to systematically evaluate bead-based enrichment in comparison to neat plasma and perchloric acid precipitation with neutralization (PCA-N). This involves experiments across multiple dimensions including pre-analytical variations commonly encountered in clinical studies, exploring the biological mechanisms underlying observed effects and analyzing the workflows in terms of their quantitative performance (**Figure 1A-D**).

**Figure 1.**
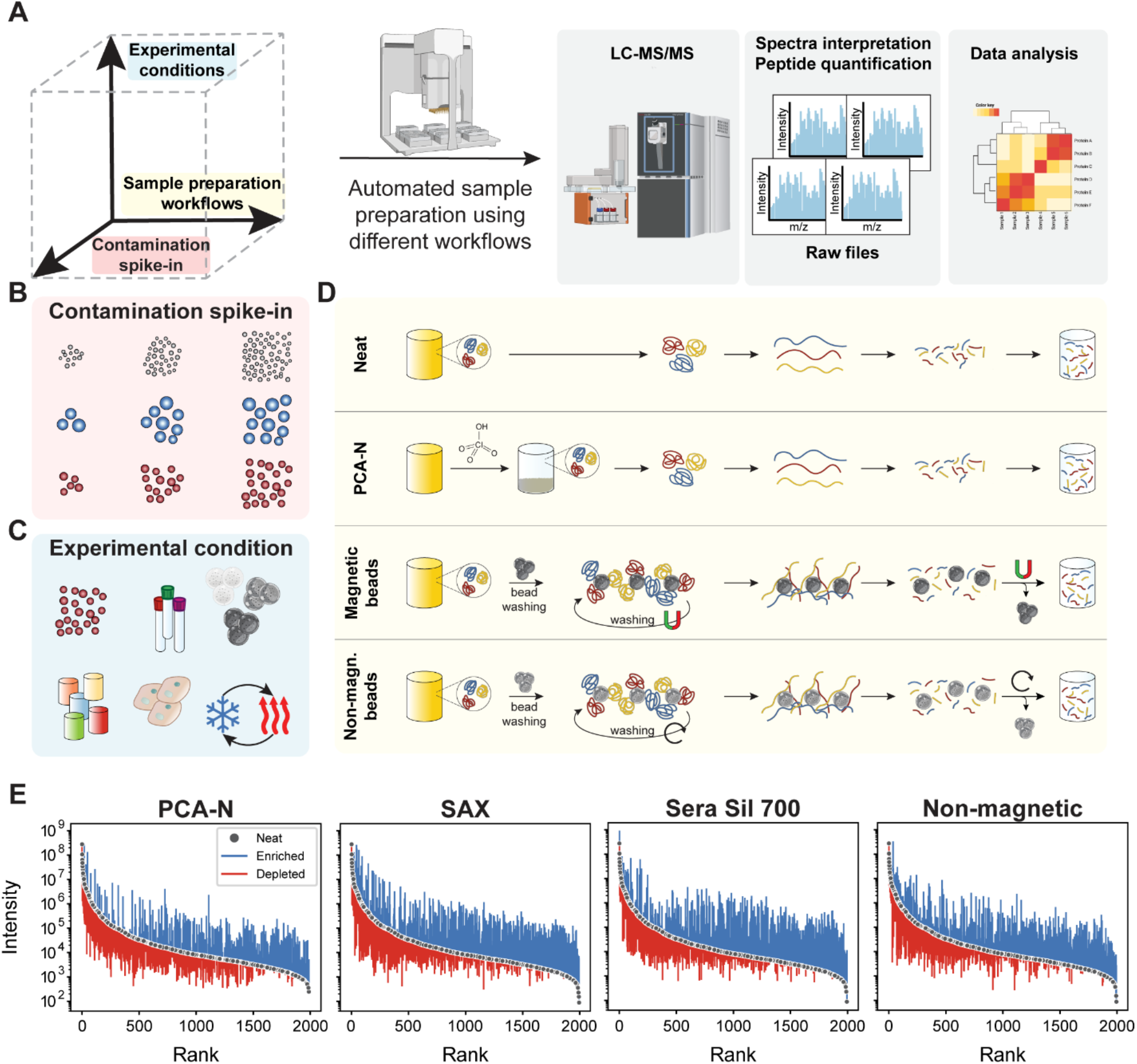
Systematic experimental framework for the comparison of plasma proteomics workflows. (A) Three-dimensional experimental design exploring workflow variations, experimental conditions, and contamination spike-in experiments. (B) Spike-in experiments consisted of defined concentration steps of platelets, erythrocytes, and PBMCs. (C) Experimental conditions include different bead types, incubation buffer compositions, and pre-analytical variations such as freeze-thaw cycles. (D) Depiction of different workflows, including sample preparation of neat plasma, perchloric acid precipitation with neutralization, and bead-based methods using magnetic and non-magnetic beads, all processed with standardized downstream analysis. (E) Rank abundance plots comparing protein intensities across workflows, showing how each method reshapes the plasma proteome by enriching low-abundance proteins (blue) and depleting high-abundance proteins (red) relative to neat plasma preparation.

We implemented five distinct plasma proteomics workflows, all automated on the Agilent Bravo Liquid Handling Platform to ensure reproducibility and high-throughput processing across 96 samples in parallel (**Methods**). On the LC-MS side we employed a state of the art workflow by coupling the Evosep chromatography system to the Orbitrap Astral (Stewart *et al*, 2023; Guzman *et al*, 2024). These workflows are meant to be representative of different analytical approaches with distinct trade-offs between throughput, depth, and susceptibility to sample quality issues. The neat plasma workflow serves as the baseline method, employing simple protein denaturation followed by enzymatic digestion, optimized for speed and simplicity. For deeper proteome coverage without beads, we evaluated perchloric acid precipitation with neutralization (PCA-N), which separates proteins based on their solubility and has shown resistance to certain contaminants. Two of our workflows utilize magnetic bead-based enrichment approaches – Strong Anion Exchange (SAX) and SeraSil-Mag 700 silica-coated superparamagnetic beads (Sera Sil 700) (**Methods**). Proteins bind to these magnetic beads, followed by on-bead denaturation and digestion, with peptides eluting during the digestion process. The fifth workflow uses non-magnetic beads with different binding characteristics and follows similar enrichment principles.

The figure also depicts our design for controlled spike-in experiments with key cellular contaminants—platelets, erythrocytes, and PBMCs—as well as the yeast proteome as a proxy for low-abundance proteins. This includes a range of analytical and pre-analytical conditions, different bead types and buffer systems, freeze-thaw cycles, centrifugation settings, blood collection tubes, and anticoagulants. This comprehensive setup enabled us to quantify each workflow’s resilience to contamination and its ability to detect low-abundance proteins under clinically relevant scenarios.

To understand how each workflow alters the original plasma proteome composition by its characteristic selective enrichment and depletion patterns, we compared changes in abundance rank orders (**Figure 1E**). Using the neat plasma workflow as our baseline, we ranked all proteins detected in neat plasma by their abundance and then tracked how each protein’s signal intensity changes in the alternative workflows. This revealed that all workflows substantially reshape the plasma proteome by enriching many low-abundance proteins while depleting a subset of high-abundance proteins. For all of the methods, this bidirectional modulation alleviates the extreme dynamic range that typically challenges comprehensive plasma proteome analysis. The PCA-N workflow is characterized by a moderate reshaping of the plasma proteome, with consistent depletion across the abundance range, allowing detection of lower abundance species. In contrast, bead-based methods lead to more dramatic transformations. The SAX and Sera Sil 700 magnetic beads achieve substantial enhancement of previously low-signal proteins, while the non-magnetic bead workflow produces the most pronounced redistribution. These differential enrichment patterns directly determine which proteins can be reliably detected and quantified in plasma samples and provide guidance for workflow selection based on target protein characteristics and research objectives.

### Contamination analysis

Building on our previous efforts to systematically evaluate the role of cellular contamination in plasma samples (Geyer *et al*, 2019), we investigated how blood cells impacted plasma proteomics workflows. Next to erythrocytes (natural range 4-6×10^6^ cells/μL of blood; ∼90 fL per cell) and platelets (1.5-4.5×10^5^ cells/μL, ∼10 fL) which are the most abundant blood cells, we added PBMCs (0.3-5×10^3^ cells/μL, ∼250 fL) (**Figure 2**). After isolating each cell type from whole blood, we verified their concentrations through complete blood counting (**Figure 2A**). Additionally, we generated pure plasma with practically no cellular contamination. We then created precise contamination steps by adding defined cell numbers to ultra-pure plasma (**Figure 2B**). These dilution series ranged from single cells up to 2×10^6^ cells. All samples were processed in quadruplicates for the five workflows, totaling 640 samples for analysis.

**Figure 2.**
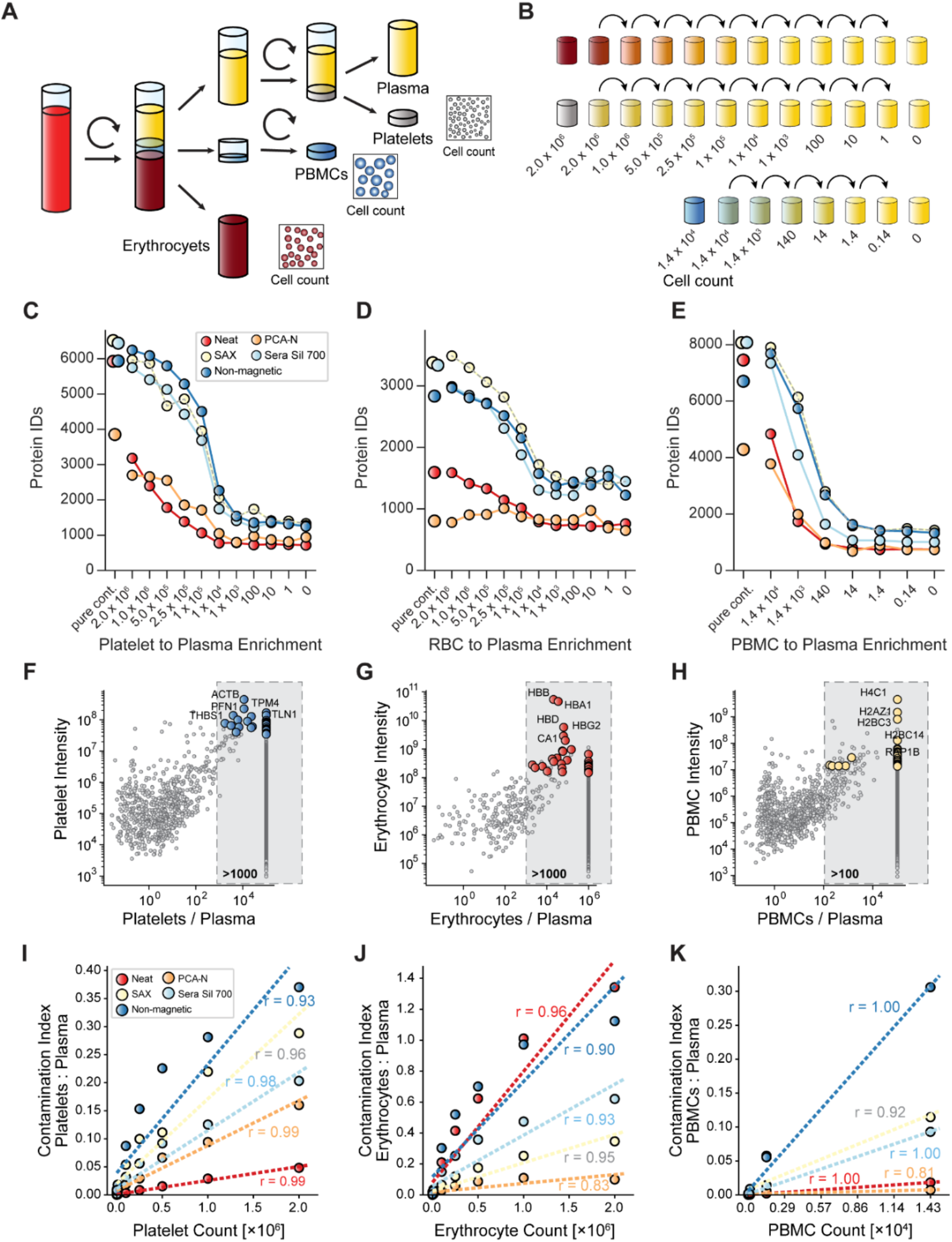
Systematic analysis of cellular contamination effects on plasma proteomics workflows. (A) Experimental design for isolation of platelets, erythrocytes, and PBMCs from whole blood. (B) Contamination series design showing concentration gradients used for spike-in experiments (1 to 2.0×10^6^ cells/µL for platelets and erythrocytes; 1 to 1.4×10^4^ cells/µL for PBMCs). (C-E) Protein identifications across contamination series in five workflows plotted against cell concentrations for (C) platelets, (D) erythrocytes, and (E) PBMCs. Data shows mean of four replicates with standard error bars. (F-H) Identification of cell-specific quality markers depicting protein intensity in pure contamination versus enrichment compared to plasma for (F) platelets, (G) erythrocytes, and (H) PBMCs. Selection criteria: >1,000-fold enrichment (platelets/erythrocytes) or >100-fold (PBMCs); minimum of two peptides (precursors) per protein; high abundance (log10 intensity >7.533 for platelets, >8.171 for erythrocytes, >7.1 for PBMCs); good reproducibility (CV <20% for platelets/erythrocytes, <35% for PBMCs). Proteins only identified in the cellular proteome were aligned to the data with a slightly higher fold change. Top five markers are highlighted. (I-K) Contamination index versus cell count for (I) platelets, (J) erythrocytes, and (K) PBMCs across all workflows. Regression lines with correlation coefficients are shown.

Analysis of protein identifications across contamination series revealed workflow-specific susceptibility patterns (**Figure 2C**). Remarkably, when spiking 14,000 PBMCs per μL of plasma, we consistently identified more than 7,000 proteins in the three bead protocols, not far from the >8,000 proteins identified in pure PBMCs. Similarly, 2×10^6^ platelets per μL plasma resulted in about 6,000 platelet proteins and 2×10^6^ erythrocytes resulted in about 3,000 proteins. Next, we investigated diminishing platelet concentrations, which revealed that bead-based workflows were exceptionally susceptible, with non-magnetic beads maintaining more than 4,500 protein identifications even at moderate contamination of 1×10^5^ platelets. This contrasted sharply with the neat workflow, where we identified 2,100 proteins when spiking in 1×10^6^ platelets/μL, roughly a doubling compared to no spike-in. PCA-N occupied a middle ground, at ∼2,600 proteins across high contamination levels but dropping to baseline at lower concentrations (**Figure 2C**).

Erythrocyte contamination, however, produced fundamentally different patterns across workflows. PCA-N demonstrated resistance to erythrocyte contamination, maintaining nearly identical protein numbers (∼800-900) regardless of spiked-in erythrocyte concentration, a unique property not observed with other cell types or workflows, which all suffered strongly from erythrocyte contamination. Despite their lower absolute numbers in the contamination series, PBMCs produced the most dramatic effects per cell (**Figure 2D**). All bead-based methods were highly susceptible to PBMC contamination down to 140 cells/μL - a full order of magnitude lower than for platelets and erythrocytes. This enhanced detection sensitivity likely reflected PBMCs’ larger size and higher protein content compared to other blood cells. When examining combined contamination effects with intermediate levels (approximately 500,000 erythrocytes/μL, 50,000 platelets/μL, and 500 PBMCs/μL), bead-based methods identified up to 8,000 proteins compared to 3,000 with the neat workflow. Collectively, these results demonstrate that blood cell contamination has the potential to substantially impact protein identification in a workflow-dependent manner, with bead-based methods showing high susceptibility to cellular protein contaminations, while PCA-N exhibits unique resistance to erythrocyte contamination. Given this potential for misleading results, we next set out to systematically assess plasma quality in measured samples by establishing robust marker panels for each cell type that could serve as indicators of contamination (Geyer *et al*, 2019).

We first compared protein abundances between pure contamination in neat plasma (**Figure 2D-F**). To calculate fold-change ratios we applied stringent filtering criteria, including >1,000-fold enrichment for platelets and erythrocytes and >100-fold for PBMCs, minimum precursor requirements, abundance thresholds, and reproducibility standards. This approach identified specific panels that included key proteins such as ACTB, PFN1, THBS1, TPM4, and TLN1 for platelets; HBB, HBA1, HBD, HBG2, and CA1 for erythrocytes; and H4C1, H2AZ1, H2BC3, H2BC14, and RAP1B for PBMCs. In characteristic distribution patterns, enrichment factors for platelet and erythrocyte markers exceeded 10^4^ (**Figure 2F,G**). Compared to our previously published markers, 20 of the topmost stringent 30 proteins were identical, confirming consistency across different sample preparation workflows and different MS instruments, scan modes and software. This demonstrated robustness of these cell-specific markers for quality assessment purposes (**Supplementary Figure 1**).

Next, we followed these quality panels across the various workflows using the dilution series experiment (**Supplementary Figure 2**). Bead-based workflows again showed high susceptibility to platelet contamination compared to the neat workflow whereas the PCA-N workflow was little affected by erythrocyte proteins. Again, PBMC markers were detectable at much lower cell counts (140 cells) compared to platelet and erythrocyte ones (10^4^ cells).

To quantify the overall impact of cellular contamination on plasma proteomics workflows as a simple metric, we calculated a contamination index for each sample by dividing the summed MS intensity of the quality markers by the summed MS intensity of all other quantified proteins (**Figure 2I-K**). For platelets all workflows showed a strong correlation between cell count and contamination index, with distinctly different sensitivity profiles (**Figure 2I**). As higher cell counts should directly result in higher contamination indices, fitting the data with a linear model worked well, with non-magnetic beads demonstrating the highest contamination index and steepest slope, followed by the magnetic bead-based workflows. The PCA-N workflow showed less substantial sensitivity to high platelet counts, while the neat workflow had a nearly flat slope, indicating minimal susceptibility to platelet contamination. This was in contrast to erythrocyte results, where neat workflow showed the highest contamination index and steepest slope, followed by the non-magnetic beads (**Figure 2J**). The magnetic bead workflows showed moderate sensitivity, while the PCA-N workflow was nearly unperturbed by erythrocyte contamination.

The contamination indices were approximately three times higher for erythrocytes than for platelets, partially due to some dominant proteins in the erythrocyte proteome and the relatively larger size of these cells. PBMC contamination (**Figure 2K**) produced patterns similar to platelets, with non-magnetic beads showing the highest sensitivity, followed by magnetic beads, while both neat and PCA-N workflows showed minimal response. We found that bead-based workflows were better described by a power law model particularly for platelet and erythrocyte contamination in bead-based workflows, suggesting saturation effects at higher contamination levels across all cell types, presumably reflecting the finite binding capacity of the beads (**Supplementary Figure 3**).

To determine a possible disproportionate contribution to contamination by cellular components, we defined a cellular enrichment score as the ratio of the summed MS intensity of the top 30 cell-specific markers by the summed intensity of the top 30 plasma proteins. For platelet contamination, non-magnetic beads had high enrichment scores of 2.5 compared to less than 0.1 for the neat workflow (**Supplementary Figure 4**). The scores for SAX and Sera Sil 700 were 1.0 and 0.5, respectively. Erythrocyte enrichment scores were more uniform across workflows, with all methods, including neat, showing similar concentration-dependent patterns. For PBMCs, non-magnetic beads again showed the highest enrichment scores while neat and PCA-N displayed minimal enrichment despite the disproportionate impact of these cells at low counts.

Of note, using spike-in experiments with contamination levels that may occur in collected samples (e.g., 5×10^5^ erythrocytes, 1×10^4^ platelets, 1,400 PBMCs), we identified approximately 8,000 proteins with bead methods compared to 3,000 with the neat workflow, while in worst-case high contamination scenarios, this number exceeded 10,000 proteins with bead-based methods.

These findings underscore our hypothesis that plasma contamination may distorts proteome profiles in a workflow and cell-type specific manner. Bead-based methods are highly sensitive, especially to platelet contamination, while PCA-N offers unique resistance to erythrocyte-derived proteins.

### Yeast spike-in as an external standard for quantitative workflow assessment

To complement the cellular contamination analysis, we next assessed each workflow’s ability to detect low-abundance proteins. For this, we spiked-in the complete soluble yeast proteome as an external standard. This allowed us to systematically evaluate enrichment efficiency and dynamic range compression across workflows, without the complication of accessing cellular vs. plasma origin of measured proteins as is the case for human proteins.

Soluble yeast proteins obtained through cell disruption and centrifugation were spiked into plasma at 8 different ratios for the 5 workflows in quadruplicates (1:2, 1:4, 1:10, 1:10^2^, 1:10^3^, 1:10^4^, 1:10^5^, 1:10^6^, and pure plasma) **(Methods**) (**Figure 3A**). In the pure yeast sample, we identified more than 4,000 proteins over 5 orders of magnitude of MS signal in our standard 100 SPD method, nearly the entire proteome (de Godoy *et al*, 2008) (**Figure 3C**). This gradually declined as a function of dilution in plasma, with the number of identified yeast proteins a proxy for the ability of the workflow to detect low-abundance proteins. In fact, we observed substantial differences between workflows (**Figure 3B**). For instance, in the 1:100 dilution ratio, the bead-based workflows demonstrated superior performance: Of the total detected 2,286 yeast proteins more than 70% (1,676) were exclusive to bead-based workflows, with 44% of these consistently identified across all three methods. The non-magnetic bead workflow performed best, uniquely identifying 282 proteins (12% of total). In contrast, PCA-N and neat workflows were less effective, yielding 471 and 317 protein groups, respectively, with minimal unique identifications. Protein identification efficiency showed clear concentration dependence, with the percentage of proteins common across all workflows decreasing from 17% at a 1:2 ratio to 9% at a 1:1,000 ratio (**Supplementary Figure 5**).

**Figure 3.**
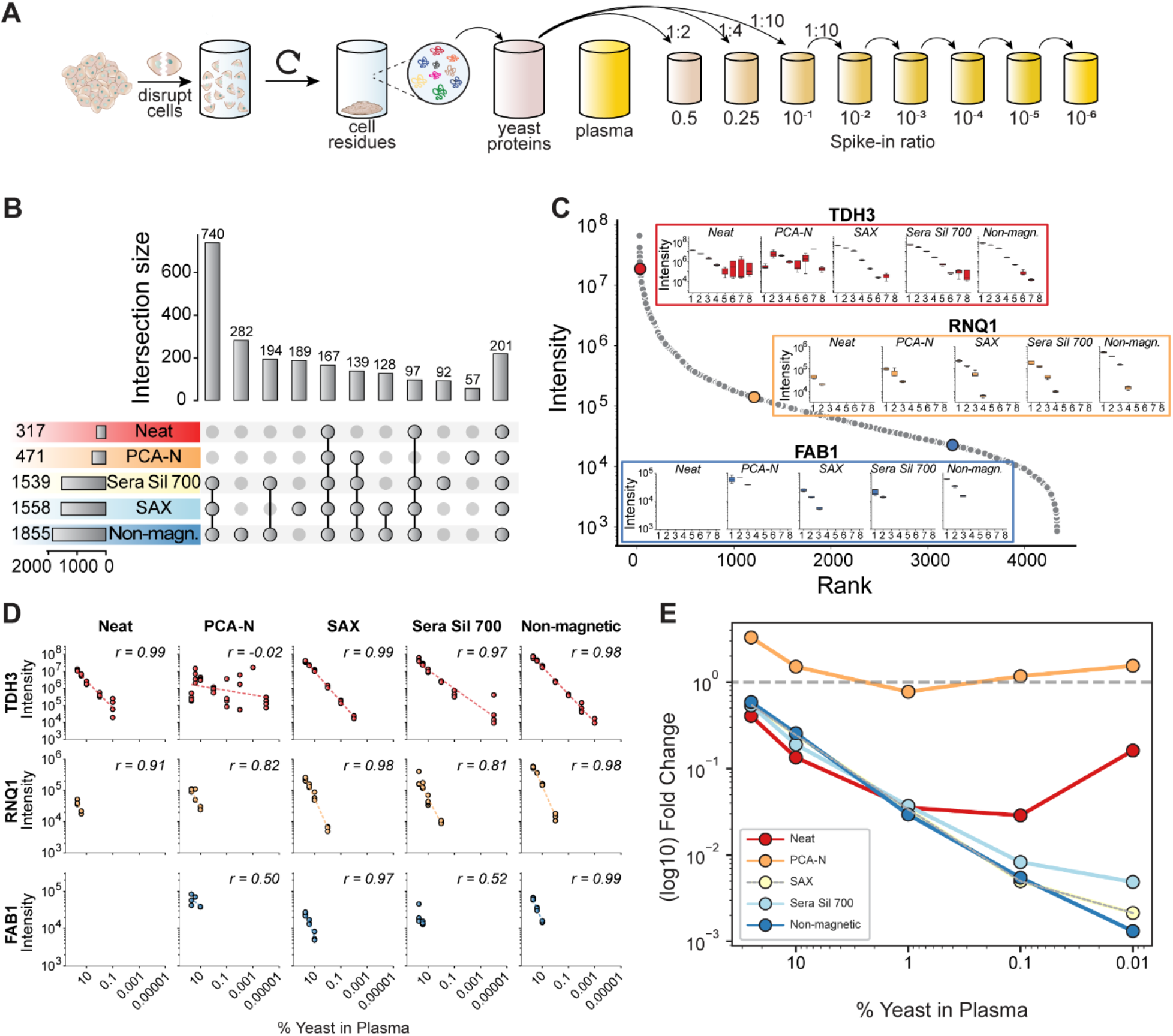
Analysis of yeast protein spike-in across plasma proteomics workflows. (A) Experimental design for yeast protein preparation and plasma spike-in series. (B) Identified yeast proteins for a 1:100 ratio, visualized using an UpSet plot. (C) Dilution series of yeast proteins, presenting abundance-ranks of three representative proteins (TDH3, RNQ1, FAB1). For this analysis, DIA-NN search was performed with 0.1% false discovery rate and data was filtered to include only proteins with at least three precursor peptides. The numbers 1-8 on the x-axis represent the dilution series with 1=1:2, 2=1:4, 3=1:10, 4=1:10^2^, 5=1:10^3^, 6=1:10^4^, 7=1:10^5^, and 8=1:10^6^ yeast:plasma ratio. (D) Workflow correlation between spike-ins, showing the relationship of protein abundances across different dilution ratios. The same data as for (C) was used, and false positives were removed for correlation analysis. (E) Signal intensities across decreasing yeast concentration. Percentage of yeast in plasma versus log10 fold change are shown from 50% yeast reference sample for the top 100 yeast proteins across all workflows.

We next evaluated the workflows for high, medium and low abundance proteins (TDH3, RNQ1 and FAB1). The highly abundant protein TDH3 was detectable in the neat workflow down to 1:10^3^ dilution, while magnetic bead-based methods extended detection by an additional order of magnitude, and the non-magnetic bead workflow by two orders of magnitude (**Figure 3C,D**). The PCA-N workflow maintained consistent median intensities but high variability across all dilution ratios. The bead workflows had high Pearson correlation coefficients of TDH3 intensity across yeast spike-in ratio (r>0.97), although not for PCA-N. Five other high-abundant proteins confirmed this pattern (**Supplementary Figure 6**). The medium-abundant protein RNQ1 was detected down to 1:4 dilution in the neat workflow, which increased to 1:10^3^ with SAX and non-magnetic beads, showing superior correlation (**Figure 3C,D**). The low-abundant protein FAB1 was not detectable in the neat workflow, but was found down to a 1:10 spike-in ratio in PCA-N and the bead-based workflows (**Figure 3C,D, Supplementary Figure 6**).

For the top 100 most abundant yeast proteins, the neat workflow showed the expected quantitative behavior down to a 1% spike-in ratio (**Figure 3E**). In the bead-based methods, intensity decreased less steeply than expected from dilution alone, enabling quantification down to 0.01% yeast in plasma - a 100-fold sensitivity improvement over the neat workflow. The observed fold-changes in PCA-N remained nearly constant across dilutions **Supplementary Figure 7**).

## Platelet activation

To shed more light on the pronounced sensitivity of bead-based protocols to platelet contamination, we designed an in-depth experiment with eight different types of beads versus seven buffer conditions. Surface chemistries ranged from negatively charged Sera Sil 700 (silanol hydroxyl groups) to positively charged SAX beads for strong anion exchange, Lewis acid-based TiO_2_ and ZrO_2_, chelating Ti-IMAC and Zr-IMAC NTA, and the non-magnetic beads. The binding milieus encompassed physiological conditions (BTP/NaCl, HEPES, PBS), modified PBS variants (EDTA/PBS, PBSC) as well as conditions known to activate platelets (CaCl_2_ and low pH environment of 2% TFA, pH 0.5). We tested these 56 conditions in quadruplicates on pure plasma containing platelets to generate a defined contaminated sample.

We first focused on total protein identifications in pure and platelet-contaminated plasma across all conditions, which varied between 650 and 2,500 in pure and between 2,000 and 3,200 in platelet-contaminated plasma (**Figure 4A,B**). Non-magnetic beads and HEPES buffer consistently yielded the highest protein identifications in pure plasma. Trends generally remained similar in platelet-contaminated plasma with notable exceptions such as generally high identifications when employing TFA or PBSC.

**Figure 4.**
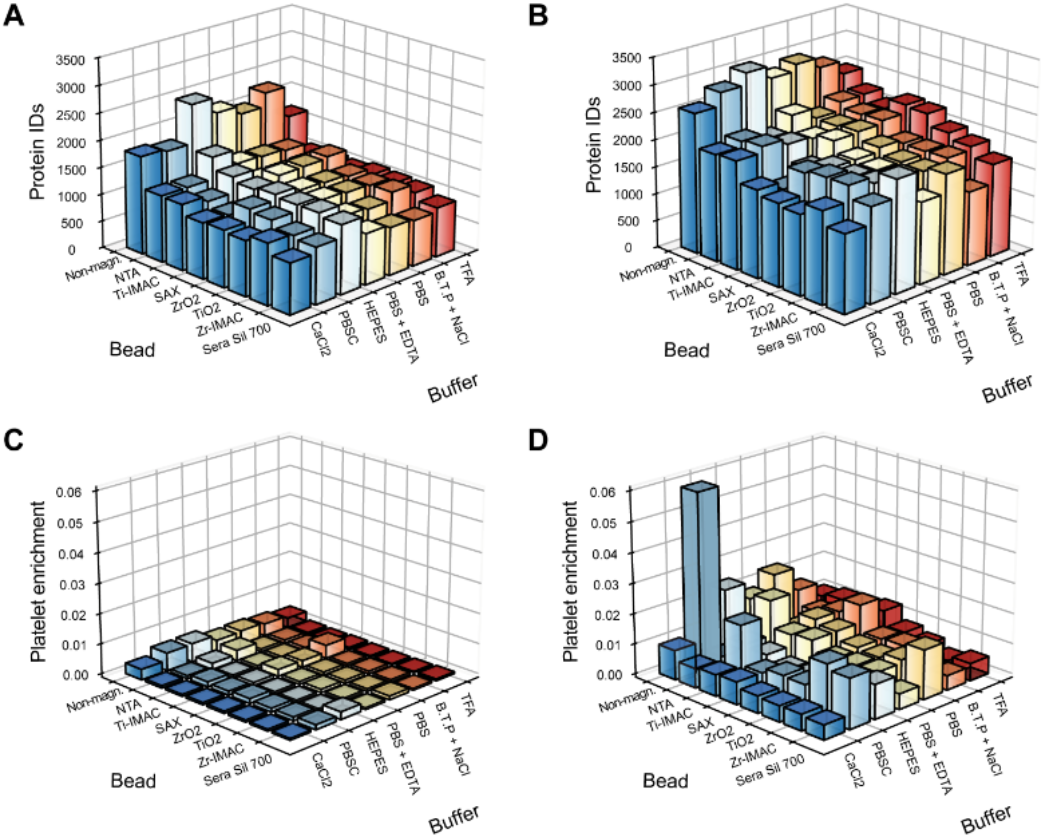
Analysis of bead and buffer combinations in pure and platelet-contaminated plasma. (A) Number of protein identifications for bead-buffer combinations in pure plasma. (B) Number of protein identifications in platelet-contaminated plasma. (C) Platelet enrichment factor across bead-buffer combinations in pure plasma. (D) Platelet enrichment factor across bead-buffer combinations in platelet-contaminated plasma.

To elucidate mechanisms behind these variations in protein numbers, we calculated the platelet enrichment factor for each condition. As expected, pure plasma samples in general showed very low platelet enrichment compared to platelet contaminated plasma (**Figure 4C,D**). However, non-magnetic beads consistently had stronger platelet enrichment across all buffers. Note that non-magnetic bead workflows required an additional centrifugation step for bead washing, which may have co-pelleted residual platelets. This likely contributed to the higher protein numbers observed with non-magnetic beads, especially given our observation that even minimal contamination (∼10^4^ platelets/μL) markedly affected this workflow. This interpretation was supported by plasma with defined platelet spike-in, where non-magnetic beads and PBSC buffer resulted in high platelet enrichment compared to the other conditions (2 to 3-fold when combined, **Figure 4D**).

Physiologically, platelets are activated during injury, leading to protein secretion, but this process can also be triggered during plasma sample preparation. We evaluated the effects of different beads and buffers on platelet activation focusing on five defined platelet markers (**Supplementary Figure 8**). Hierarchical clustering revealed that PF4V1 and PPBP had low intensities when employing CaCl_2_, TFA buffers or Sera Sil 700 beads, which have negative surface charge. These conditions activated platelets, leading to release of these proteins (Singh *et al*, 2019). In contrast, non-magnetic beads with PBSC buffer led to high intensities for the activation markers, indicating platelet enrichment without activation.

### Rescue of platelet-contaminated samples

Given that already collected plasma samples frequently have significant platelet contamination, which is readily apparent in sensitive plasma proteomics workflows, especially bead-based ones, we asked to what extent this could be mitigated or ‘rescued’. To this end, we investigated defined platelet spike-ins, which we subjected to either direct processing or repeated freeze-thaw cycles (3x -80°C for 15 min, thawing at 37°C for 10 min). After splitting, these samples underwent centrifugation (3000g, 30 min) with careful supernatant collection, or no centrifugation. The resulting eight conditions were analyzed in quadruplicate using all five workflows (**Figure 5A**).

**Figure 5.**
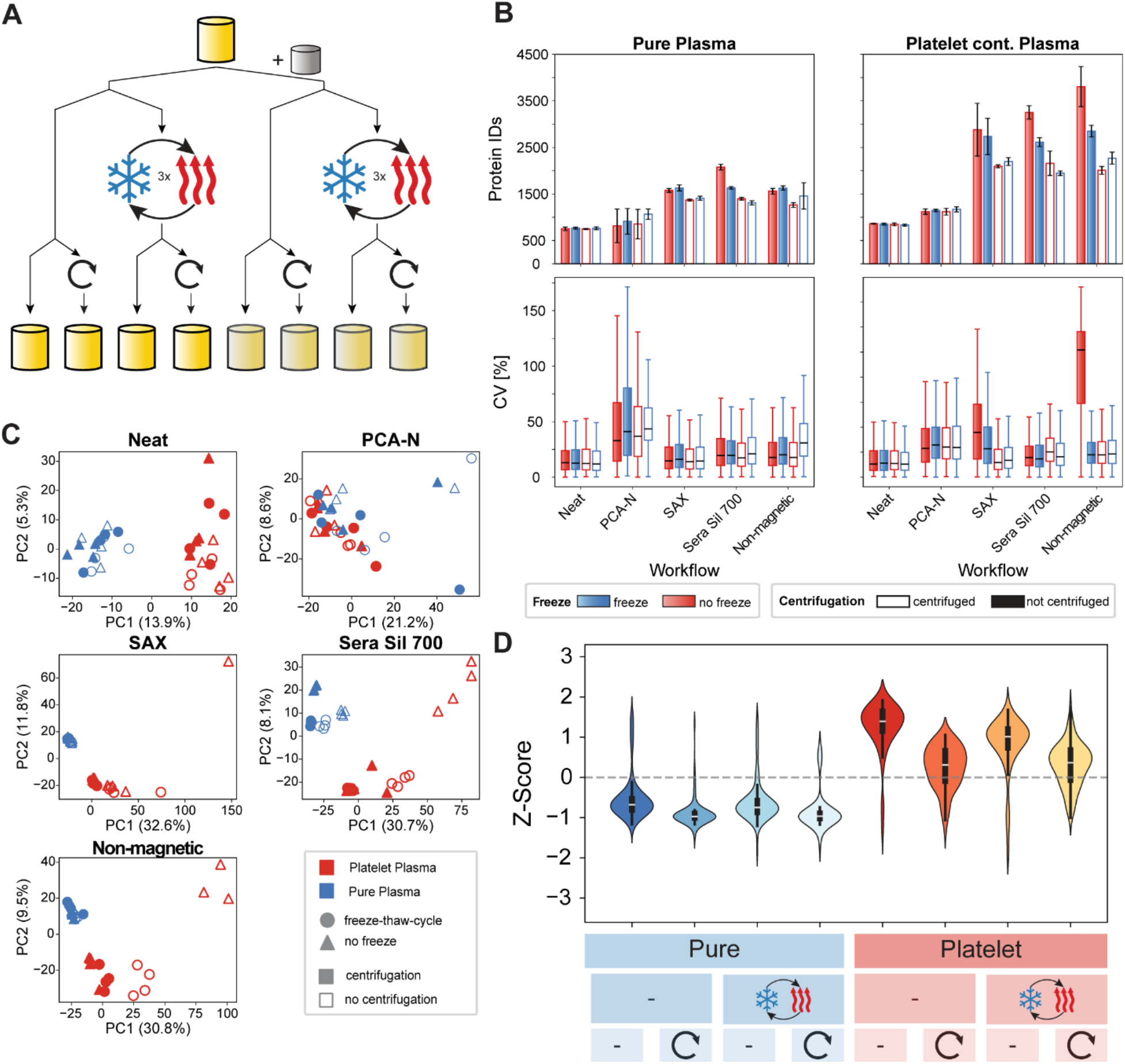
Assessment of plasma processing strategies for platelet contamination rescue. (A) Experimental design for plasma sample processing evaluation. (B) Number of protein identifications (top) and CV distributions (bottom) for pure and platelet-contaminated plasma across five workflows. (C) Principal component analysis comparing different processing conditions. Percentage variance explained by PC1 shown in parentheses. One outlier (Platelet-contaminated plasma without freeze-thaw cycle and not centrifuged, replicate 1, Non-magnetic beads) was removed from the analysis, and K-nearest neighbors (KNN) imputation with 3 neighbors was applied to maintain dataset completeness. (D) Violin plots showing Z-scored intensities of the top 100 platelet markers across all conditions and workflows. Blue represents pure plasma samples, red shows platelet-contaminated samples.

The neat workflow was remarkably stable across all conditions with only minor variations in identifications and quantification (∼800 +/- 50 proteins, median CV ∼13%), followed by PCA-N with a 35% increase in identifications. Identifications in magnetic bead-based workflows increased from 1,400 proteins in pure plasma to 2,700 proteins in platelet-contaminated samples, which decreased by only 15-20% after centrifugation (**Figure 5B**). For non-magnetic beads the reduction was larger, from 3,700 to about 2,000.

A Principal Component Analysis (PCA) pointed to platelet contamination as the dominant source of variation, whereas the contribution of freeze-thaw cycles was minimal (**Figure 5C**). In all cases except PCA-N, the PCA clearly separated centrifuged and non-centrifuged samples in the rescue experiments, supporting the benefit of this step. This was also apparent in violin plots showing Z-scored intensities of the top 100 platelet markers across all conditions (**Figure 5D**). In pure plasma samples, intensities of these remained consistently low and stable across all processing conditions. In contrast, without centrifugation platelet-contaminated samples retained high platelet marker intensities regardless of freeze-thawing. However, centrifugation still substantially reduced platelet marker intensity in contaminated samples. This reduction suggested partial rescue of platelet-contaminated samples. This pattern was consistent across all workflows, with bead-based methods showing the most pronounced rescue effect as was also evident at the single quality marker level (**Supplementary Figure 9**).

## Recommended plasma sample preparation conditions

While post-collection interventions such as centrifugation could reduce the impact of contamination, they did not universally restore sample quality. To identify optimal sample handling conditions, we systematically evaluated how pre-analytical variables, particularly centrifugation settings and tube types, affected contamination levels and downstream proteome profiles. We collected blood from 11 healthy individuals into standard EDTA tubes and EDTA gel separator tubes and subjected each to four protocols: 500g for 7 min, 1,000g for 7 min, 3,000g for 7 min, and 3,000g for 30 min (**Figure 6A**). Clinical blood counts revealed that gel tubes retained substantially more cellular material at low speeds, likely because the gel barrier activated only at higher g-forces. At 500g, gel tubes contained approximately 570,000 platelets per μL compared to 320,000 in standard tubes - a nearly 2-fold enrichment compared to whole blood. Erythrocyte contamination followed a similar trend, with higher levels in gel tubes under mild centrifugation. Increasing speed and time dramatically reduced contamination: platelet counts dropped below 6×10^4^/µL at 3,000g for 7 min and became undetectable after 30 min. Erythrocytes were effectively removed at 3,000g in both tube types, and PBMCs decreased from ∼670/µL in gel tubes at 500g to near-zero across higher-speed conditions. With these contamination profiles established, we processed all samples through our five distinct workflows and analyzed them by mass spectrometry. Protein identification numbers were strongly dependent on both centrifugation conditions and the chosen workflow (**Figure 6B**). Bead-based enrichment identified substantially more proteins than neat or PCA-N at lower speeds. At 500g, all three bead workflows identified over 5,500 proteins, whereas the neat and PCA-N workflows yielded only about 2,000 proteins. The effect of increasing centrifugation was most dramatic for bead-based methods where protein identifications dropped sharply with g-force. For instance, the number of identified proteins in the non-magnetic workflow decreased by approximately 70% between the lowest and highest centrifugation conditions. When comparing gel and non-gel tube types, they resulted in comparable protein identifications, with differences becoming apparent only at extended centrifugation times (**Figure 6C,D**).

**Figure 6.**
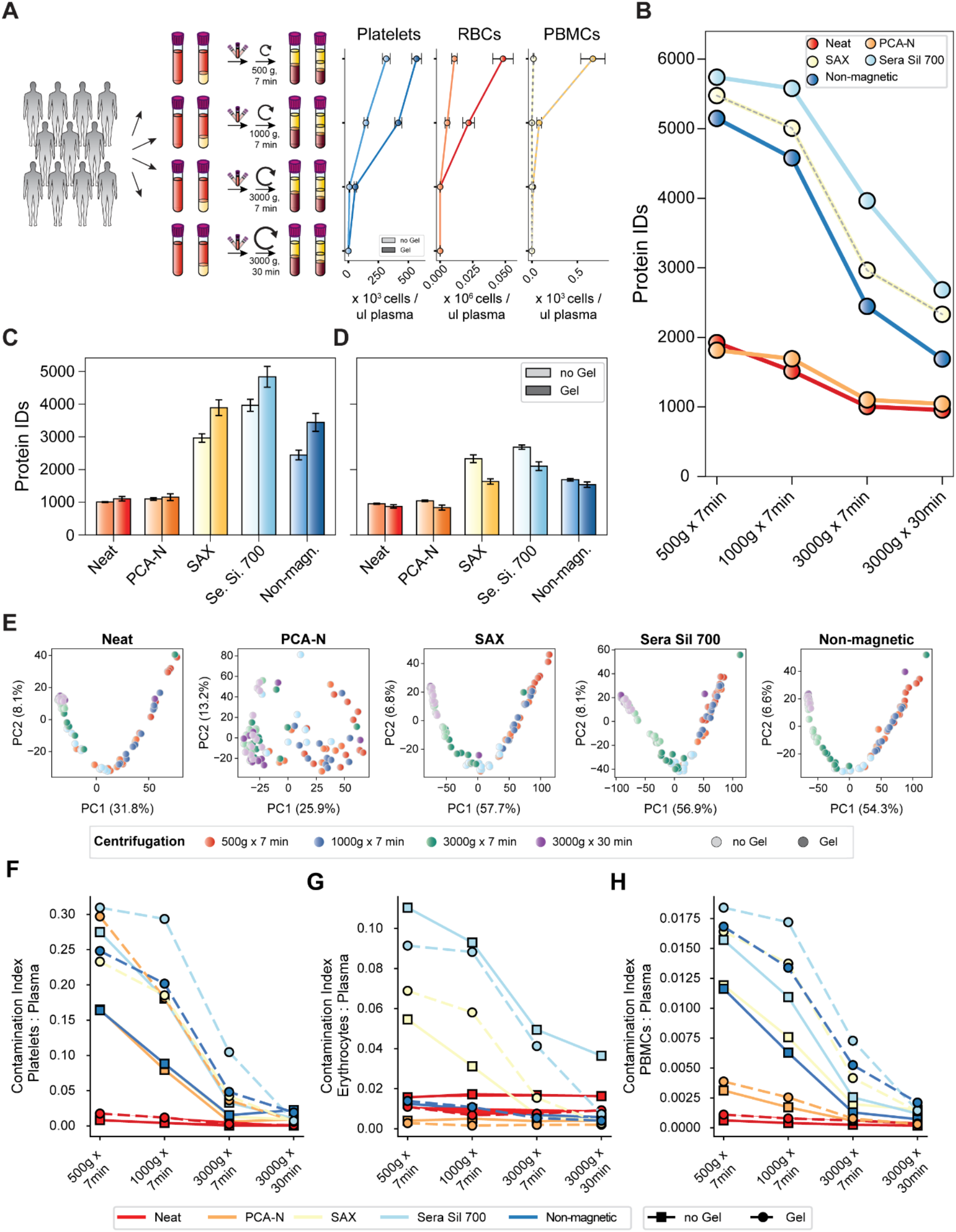
Impact of centrifugation conditions on plasma proteome analysis. (A) Experimental design: Blood was collected from 11 healthy individuals into standard EDTA tubes and EDTA gel separator tubes, followed by centrifugation at four different conditions (500g for 7 min, 1,000g for 7 min, 3,000g for 7 min, and 3,000g for 30 min). Clinical measurements of cellular contamination (platelets, RBCs, PBMCs) were performed. (B) Number of protein identifications across five workflows plotted as lines, shown for standard EDTA tubes without gel separator. (C) Number of protein identifications at 3,000g for 7 min comparing standard tubes versus gel separator tubes across all five workflows. (D) Number of protein identifications at 3,000g for 30 min comparing standard tubes versus gel separator tubes across all five workflows. (E) Principal component analysis of proteomics data for each workflow, colored by centrifugation condition and tube type. (F-H) Contamination indices plotted as lines across centrifugation conditions for (F) platelets, (G) erythrocytes, and (H) PBMCs. Solid lines represent standard tubes while dashed lines represent gel separator tubes, with different colors indicating different workflows.

Principal component analysis highlighted centrifugation as the main factor shaping proteome profiles, with PC1 explaining between 26% (PCA-N) and 58% (SAX) of the variance (**Figure 6E**). Bead-based workflows clearly clustered by g-force, especially at 3,000g for 30 min. In contrast, PCA-N exhibited minimal separation, indicating low sensitivity to processing variation. The distinction between gel and no-gel tubes was most evident at lower speeds and nearly disappeared at 3,000g for 30 min. Tube type influenced clustering mainly at low speeds.

Contamination indices provided a clear quantitative assessment of residual cellular content (**Figure 6F-H**). Bead-based workflows showed markedly elevated platelet contamination indices at low centrifugation speeds, especially in gel tubes (**Figure 6F**). Platelet contamination decreased steadily with increasing g-force and time, with over 90% reduction between the lowest and highest conditions. However, even at 3,000g, bead workflows retained 10–20 times more platelet signal than the neat workflow. This shows that high centrifugation speeds are necessary to decrease platelet contamination, especially for bead-based workflows. The contamination bias of bead-based workflows completely settles in the range of the neat workflow only at long times. Nevertheless, even at very low concentrations platelets are disproportionately captured and enriched by bead surfaces.

For erythrocyte and PBMC contaminations neat and PCA-N showed low or no contamination while bead-based enrichments were highly sensitive to both contaminations, especially at low centrifugation speed (**Figure 6G,H**). PBMC contamination showed less pronounced contamination pattern than platelets, with 5-fold enrichment for bead-based methods over the neat plasma workflow.

To complement the aggregate contamination indices, we analyzed individual marker proteins across centrifugation conditions (**Supplementary Figure 10B**). PPBP, HBB, and H4C1 revealed workflow-specific enrichment patterns, underscoring the added value of single-protein analysis.

We next assessed the impact of anticoagulant type and gel separators by collecting blood into EDTA, Li-Heparin, and serum tubes, with and without gels, from the 11 donors, all processed identically (3,000g, 7 min; **Supplementary Figure 11A**). Neat and PCA-N workflows were largely unaffected by matrix type (∼1,000 proteins identified), whereas bead-based workflows showed strong variation, with the highest identifications in EDTA plasma (up to 5,000), followed by serum and Li-Heparin (**Supplementary Figure 11B)**. Principal component analysis showed clustering primarily by anticoagulant rather than gel use, most notably in the Sera Sil 700 workflow (**Supplementary Figure 11C**). Platelet and erythrocyte contamination was highest in EDTA for bead workflows, while PCA-N remained resistant across conditions (**Supplementary Figure 11D**).

Based on these findings, we recommend that anticoagulant type should be standardized within a study. Combining matrices, such as EDTA and Li-Heparin treated samples, should be strictly avoided. Although EDTA is a frequently used anticoagulant in biomarker research, our data suggests that other blood collection metrices like Li-Heparin and serum provide less contaminated samples for proteomics experiments, especially when applying bead-based workflows.

## Discussion

Our systematic comparison of five distinct plasma proteomics workflows—including three bead-based protocols (Blume *et al*, 2020; Wu *et al*, 2024), a perchloric acid-based precipitation neutralization (PCA-N) (Albrecht *et al*, 2025), and a conventional neat plasma workflow—reveals a critical trade-off between proteome depth and susceptibility to cellular contamination. Bead-based methods provide the highest protein identifications even in clean plasma but are acutely sensitive to contamination with platelets, erythrocytes, and PBMCs. In contrast, PCA-N shows strong resistance, especially to erythrocytes and, to a lesser extent, platelets. The neat workflow occupies a middle ground with moderate vulnerability but fewer identifications. The latter two technologies offer advantages of automation, low additional costs, and ease of use.

Using soluble yeast proteins as an orthogonal spike-in strategy, we confirmed that bead-based methods quantitatively enrich low-abundance proteins beyond the capabilities of neat or PCA-N workflows. This enrichment was accurate across several orders of magnitude and confirmed dynamic range compression in bead-based approaches. PCA-N featured consistent protein detection across abundance categories despite higher variability, suggesting selective protein depletion and enrichment mechanisms that enable low-abundance protein detection while resisting certain contamination types—a behavior also noted by Beimers et al. (2025) in their comparison of acid-based depletion methods. Thus, the increased sensitivity is real—but comes at the cost of increased vulnerability to bias when working with non-ideal specimens.

This vulnerability has profound implications for biomarker discovery. In case-control studies, systematic differences in sample quality between groups can mimic or obscure true biological effects, particularly when using bead-based workflows. A small variation in platelet contamination may manifest as hundreds or thousands of additional proteins, compromising statistical power and potentially leading to spurious biomarker candidates. This calls into question the suitability of bead-enrichment methods for studies using archived or clinically collected plasma samples unless stringent quality control measures are in place. Our findings are particularly relevant for longitudinal studies where sample collection may span months or years with potential variations in handling protocols, and for cross-sectional and multicentric studies comparing different patient populations where sample collection conditions might systematically differ between sites.

To mitigate these risks, we have previously developed a three-step contamination control strategy: (1) assessing contamination in individual samples using our validated marker panels, (2) detecting potential bias between study groups, and (3) correlating candidate biomarkers with cell-specific markers to identify artifacts. Building on our previous work (Geyer *et al*, 2019), we developed contamination indices and validated quality panels for platelets, erythrocytes, and PBMCs. These were consistent across workflows and instruments and enable a reliable assessment of contamination at both sample and cohort levels. Notably, in the presence of contaminating cells, bead-based workflows result in an inflation of protein identification. While PBMCs occur at much lower concentrations than erythrocytes or platelets in blood, they produced substantial proteome alterations at just 140 cells/μL in bead-based workflows, highlighting the critical importance of controlling for cellular contamination in studies examining immune-related conditions (Robinson *et al*, 2013; Messner *et al*, 2020).

Post-collection interventions—particularly centrifugation at 3,000g for 30 min after thawing— can partially rescue compromised samples by reducing platelet marker signals, though not entirely eliminating them. This finding addresses a critical gap in current plasma proteomics guidelines, which typically recommend optimal collection procedures but provide limited guidance for handling already-compromised samples (Ignjatovic *et al*, 2019). We therefore recommend this centrifugation protocol as a standard step in any proteomic analysis of stored plasma.

Beyond rescue strategies, we systematically evaluated centrifugation conditions and anticoagulant types as pre-analytical variables. Low-speed centrifugation or the use of gel separator tubes were associated with high residual cellular content, particularly platelets. These effects were workflow-dependent: while neat and PCA-N workflows remained relatively unaffected, bead-based workflows were highly sensitive. Even after high-speed centrifugation, residual platelet signals remained detectable in bead-based methods—suggesting that platelets are selectively retained or enriched by certain bead surfaces.

Our detailed examination of centrifugation conditions and tube types revealed critical insights for plasma sample preparation. At lower forces (500g, 1000g), gel separator tubes actually increased contamination compared to standard tubes, as the gel begins to move but does not form a complete barrier, creating turbulence (Bowen & Remaley, 2014; Sadgrove *et al*, 2024). Centrifugation at 3,000g for 30 min provided the cleanest plasma across all tube types, minimizing contamination across all workflows. We therefore recommend this condition as a standard for plasma collection in proteomics studies, particularly those employing bead-based enrichments, although 3,000g and 7 min may often be more practical. Consistency in tube type and anticoagulant (e.g., using EDTA plasma rather than serum or Li-Heparin) also proved critical. EDTA plasma yielded the highest and most stable identifications in bead workflows, whereas Li-Heparin introduced distinct proteome patterns, likely due to its effect on protein binding and coagulation. The complex relationship of pre-analytical variation and detectable proteome is emphasized throughout the plasma proteomics literature (Deutsch *et al*, 2021; Geyer *et al*, 2019; Ignjatovic *et al*, 2019).

Our findings are supported by a very recent preprint from Gao et al. (2025). This thorough and well-designed study independently showed that nanoparticle-based enrichments are highly susceptible to blood cell contamination, particularly from platelets and erythrocytes. Their use of bovine plasma dilution series mirrors our yeast spike-in design and independently confirms the quantitative capability—but also the contamination risk—of bead-based methods (Gao *et al*, 2025).

Taken together, our study provides both a cautionary note and a practical framework. Bead-based enrichments offer real and powerful gains in depth, but these must be weighed against a heightened risk of systematic bias. Our three-step contamination control strategy for biomarker studies provides practical guidance: Contamination Assessment through calculating indices for each sample, Bias Detection by evaluating whether contamination markers differ between groups, and Candidate Validation by correlating biomarkers with cell-specific intensities. As MS instrumentation continues to improve, with platforms like the Orbitrap Astral pushing depth and throughput further (Stewart *et al*, 2023; Hendricks *et al*, 2024; Lancaster *et al*, 2024; Serrano *et al*, 2024), and new application areas of plasma proteomics expand (Niu *et al*, 2025), the importance of controlling for sample quality will only grow. As technological capabilities expand, there will be a growing need to refine and broaden contamination panels to include additional cell types and to further resolve PBMCs into their subpopulations.

In conclusion, our study underscores that sample quality—particularly in the form of cellular contamination—remains a key determinant of plasma proteomics data. By identifying the strengths and vulnerabilities of commonly used workflows, we offer guidance for selecting and optimizing methods based on sample context. We encourage researchers to apply rigorous contamination assessment and to match analytical depth with robustness, thereby ensuring that plasma proteomics fulfills its promise in biomarker discovery and translational research.

## Acknowledgments

We thank all members of the Proteomics and Signal Transduction Group for help and discussions, and in particular Andre C. Michaelis, Igor Paron, Tim Heymann and Katharina Zettl for technical assistance. We thank Florian Arendt, and Britta Pauli for their clinical assistance and all blood donors who made this research possible. This work is supported by the Max Planck Society for Advancement of Science.

## Author contributions

KK performed and interpreted the MS-based proteomic analysis of patient plasma; wrote the paper; and generated the figures. PEG performed together with JBMR, VA, ASS and EI experiments and generated article text. PEG and JBMR designed figures. DT and LMH designed experiments, drafted practical considerations for sample preparation, and worked on the article text. PEG and MM designed and interpreted the MS-based proteomic analysis of plasma, supervised and guided the project, and wrote the manuscript.

## Materials and Methods

### Blood collection and fractionation

If not otherwise indicated, whole blood was collected by venipuncture into EDTA-containing tubes (9 ml). The study was approved by the Ethics Committee of LMU Munich (Reg. No. 17-012). All participants have given written informed consent to participate in the study. For the isolation of blood components, the tubes were first centrifuged at 500g for 7 min to separate platelet-rich plasma from the cellular components. The supernatant (platelet-rich plasma) was carefully transferred to a new centrifugation tube and centrifuged again at 500g for 7 min. This supernatant was collected and centrifuged at 3,000g for 7 min. After this step, the supernatant was subjected to a final centrifugation at 3,000g for 7 min to obtain platelet-free plasma, with care taken to avoid the bottom approximately 500 μl to prevent platelet contamination. For platelet isolation, the pellet from the 3,000g centrifugation was resuspended in 4 ml PBS/EDTA (1.6 mg/ml EDTA), centrifuged at 3,000g for 7 min, and the supernatant was discarded. This washing step was repeated once more, resulting in purified platelets. To isolate erythrocytes, the pellet from the initial 500g centrifugation was transferred to a 15 ml tube and centrifuged at 3,000g for 7 min. The supernatant, buffy coat, and top 1 ml of erythrocytes were discarded. The remaining erythrocytes were washed twice by resuspending in 4 ml PBS/EDTA and centrifuging at 3,000g for 7 min, with removal of the supernatant and top layer of cells after each wash, yielding purified erythrocytes.

PBMCs were isolated from whole blood collected in CPT-Vacutainers. The vacutainers were gently inverted ten times immediately after collection and centrifuged at 1,650g for 20 min. After centrifugation, the content of the CPT-Vacutainers was transferred to 50 ml tubes. The tubes were filled with PBS buffer up to 50 ml and gently inverted to mix. These samples were then centrifuged at 400g for 15 min. The supernatant was carefully removed, and the cell pellet was retained. A filter was placed on a second 50 ml Falcon tube, and 10 ml of PBS buffer was added to the cell pellet. The cells were carefully resuspended and filtered into the second tube. Additional PBS buffer was added through the filter to reach the 50 ml mark. This suspension was centrifuged at 300g for 15 min. After discarding the supernatant, the cell pellet was resuspended in 1,600 μL of PBS buffer.

Cell counts were determined using a Sysmex XN 1,000/9,100 automated hematology analyzer (Sysmex Corporation) according to standardized laboratory procedures. This platform employs multiple measurement technologies optimized for specific cell types to ensure accurate quantification across diverse sample conditions. Platelet counts were assessed using a dual-methodology approach. The primary measurement utilized impedance measurement with hydrodynamic focusing, where platelets passing through a microaperture generate electrical pulses proportional to their volume. For samples with potential counting interferences or suspected thrombocytopenia, the analyzer additionally employed fluorescence flow cytometry (PLT-F), which labels platelets with a proprietary fluorescent marker for enhanced detection and discrimination from other cellular elements. Erythrocyte enumeration was similarly performed via impedance measurement with hydrodynamic focusing, with each erythrocyte generating a voltage pulse proportional to its volume as it passes through an electrically charged aperture. Total leukocyte counts were determined using fluorescence flow cytometry. This technique employs fluorescent dyes that differentially stain cellular components, allowing the analyzer to distinguish leukocytes based on their size, internal complexity, and fluorescence characteristics, providing accurate quantification of white blood cells in the samples.

### Sample preparation workflows

All sample preparation workflows were automated using an Agilent Bravo Liquid Handling Platform (Geyer *et al*, 2016).

Neat Plasma Workflow: Plasma (1 µL) was combined with 50 µL lysis buffer (100 mM Tris pH 8.0, 40 mM chloroacetamide, 10 mM TCEP). Samples were heated at 95°C with agitation for 10 min on a Thermomixer C. After cooling to room temperature, 10 µL digestion buffer (8 µL lysis buffer, 1 µL trypsin [0.5 µg/µL], 1 µL Lys-C [0.5 µg/µL]) was added and proteins were digested at 37°C for 16 hours. The digestion was terminated by the addition of an equal volume 0.2% TFA.

Perchloric Acid precipitation with neutralization (PCA-N): Plasma (5 μL) was diluted in 20 μL ddH_2_O followed by the addition of 25 μL 1 M perchloric acid. Samples were agitated at 4°C for 1 h, followed by centrifugation at 4,000g for 20 min at 4°C. The supernatant (24 μL) was collected and combined with 8 μL 1.4 M sodium hydroxide solution to adjust pH to 8-8.5. Lysis buffer (8 μL; containing 40 mM chloroacetamide, 20 mM DTT, 0.01% DDM, 60 mM TEAB) was added. Proteins were digested using trypsin/LysC (1.6 µL: 0.4 µL trypsin [0.5 µg/µL] and LysC [0.5 µg/µL each, 10.5 µL 1M TEAB, 0.7 µL water) and digestion was stopped with TFA (final concentration 0.5%) (Albrecht *et al*, 2025). The peptide mixtures were analyzed by LC-MS/MS.

Magnetic Bead-Based Enrichment (SAX, Sera Sil 700, TiO_2_, ZrO_2_, Ti-IMAC, Zr-IMAC, NTA): Magnetic beads (10 µL slurry, 20 mg/mL, MagReSyn® SAX, MagReSyn® TiO_2_, MagReSyn® ZrO_2_, MagReSyn® Ti-IMAC, MagReSyn® Zr-IMAC, MagReSyn® NTA, Resyn Biosciences, Sera Sil 700: SeraSil-Mag 700 silica coated superparamagnetic beads) were washed three times with incubation buffer (SAX: 100 mM Bis-Tris Propane, pH 6.3, 150 mM NaCl; Sera Sil 700: 1 M HEPES) using magnetic separation (Wu *et al*, 2024). Plasma (10 µL) was combined with 100 µL incubation buffer (SAX: 50 mM Bis-Tris Propane, pH 6.5, 150 mM NaCl; Sera Sil 700: 1M HEPES) and pre-washed beads, followed by incubation at 37°C for 30 min with agitation. After magnetic separation, bead-bound proteins were washed with washing buffer. Proteins were denatured, reduced, and alkylated while bound to the beads by the addition of lysis buffer (50 µL, 100 mM Tris pH 8.0, 40 mM chloroacetamide, 10 mM TCEP) followed by heating at 95°C for 10 min. Digestion proceeded as described for the neat workflow, resulting in peptide elution from the beads. The eluted peptides were acidified and analyzed by LC-MS/MS.

Non-magnetic bead workflow: The non-magnetic bead workflow utilized non-magnetic beads (OmniProt™, Westlake Omics), requiring centrifugation at 4,000g for 20 min for all washing and collection steps. Beads were resuspended and washed in PBS. Plasma protein binding was performed in PBS containing 0.05% CHAPS, followed by washing of bead-bound proteins with 33% PBS. Proteins were denatured, reduced, and alkylated while bound to the beads by adding lysis buffer (33.3 µL; 40 mM chloroacetamide, 20 mM DTT, 0.01% DDM, 60 mM TEAB). Digestion proceeded as described for the neat workflow, resulting in peptide elution from the beads. The eluted peptides were acidified and analyzed by LC-MS/MS.

### Data acquisition by mass spectrometry

Samples were analyzed using the Evosep One liquid chromatography system (Evosep) (Bache *et al*, 2018) coupled to an Orbitrap Astral mass spectrometer (Stewart *et al*, 2023; Hendricks *et al*, 2024) (Thermo Fisher Scientific). Chromatographic separation was performed on an 8 cm Aurora Rapid XT UHPLC column (AUR3-80150C18-XT, Ionopticks) at 50°C using the ‘100 samples per day’ method with pre-formed gradients and a total runtime of 11.5 min per sample. Mobile phases consisted of 0.1% formic acid in water (buffer A) and 0.1% formic acid in acetonitrile (buffer B). For each sample, 200 ng of peptides were loaded onto C-18 tips (Evotip Pure, Evosep) according to the manufacturer’s protocol. Mass spectrometric analysis was performed using a data-independent acquisition (DIA) method (Guzman *et al*, 2024). The ion source was operated with a static spray voltage of 1900 V in positive ion mode, and the ion transfer tube temperature was set to 280°C. FAIMS (high-field asymmetric waveform ion mobility spectrometry) was utilized in standard resolution mode with a compensation voltage (CV) of -40 V and a carrier gas flow of 3.5 L/min. Full MS1 scans were acquired in the Orbitrap analyzer at a resolution of 120,000 FWHM over a scan range of 380-980 m/z. The RF lens was set to 40%, and a normalized AGC target of 500% with a maximum injection time of 3 ms was used. DIA MS2 scans covered the same mass range (380-980 m/z) divided into 150 isolation windows of 4 Th each with window placement optimization enabled. MS2 spectra were acquired with an HCD collision energy of 25%, a normalized AGC target of 500%, and a maximum injection time of 7 ms. The scan range for fragment ions was set to 150-2000 m/z. Data were collected in profile mode for MS1 and centroid mode for MS2 scans. The expected chromatographic peak width was set to 5 seconds, and advanced peak determination was enabled to optimize duty cycle. Quality control samples were analyzed regularly throughout the analytical sequence to monitor system performance and stability.

### Data analysis / spectral search

Each raw file was converted to the mzML format using Thermo Fisher Scientific’s ThermoRawFileParser (v4.3) with format parameter set to 1, metadata parameter set to 0. The conversion was processed in parallel on a high-performance computing. The resulting mzML files were processed using DIA-NN (version 1.8.1) (Demichev *et al*, 2020) on a high performance computing cluster. The searches were performed against a human UniProt Swiss-Prot isoform database that included oxidation and N-terminal acetylation modifications using the DIA-NN build in in-silico library prediction. All files were analyzed with match-between-runs enabled using the “--use-quant” and “--reanalyse” parameters. The following parameters were applied: peak centering, smart profiling, retention time profiling, and relaxed protein inference. Further, mass accuracy was set to 10 ppm for both MS1 and MS2 scans and the scan window to 7. False discovery rate was controlled at 1% at the peptide-to-spectrum match level. Protein quantification was performed using MS1 and MS2 data with no interference signal removal. Quantitative matrices were generated for downstream statistical analyses.

### Bioinformatics analysis

All bioinformatic analyses were performed using Python in Jupyter notebooks. Data processing was conducted using pandas for manipulation of proteomics data matrices and NumPy for numerical operations, while visualization and statistical analyses employed using scipy, matplotlib, and seaborn packages. For protein quantification, we used the protein-level reports from DIA-NN output. Protein intensities were log10-transformed prior to further analysis. Where complete data matrices were required e.g., for principal component analysis, K-nearest neighbors imputation (n neighbors = 3) was applied to handle missing values.

Contamination analysis: We calculated contamination indices by dividing the summed intensity of cell-specific marker proteins by the summed intensity of all other quantified proteins. The relationship between cell counts and contamination indices was evaluated using both linear and non-linear regression models. For evaluation of cellular contamination, enrichment scores were calculated by dividing the summed intensity of the 30 defined cell-specific marker proteins by the summed intensity of the top 30 most abundant plasma proteins, to highlight the relative abundance of contamination markers compared to the dominant plasma proteome components.

## Supplementary Figures

**Supplementary Figure 1.**
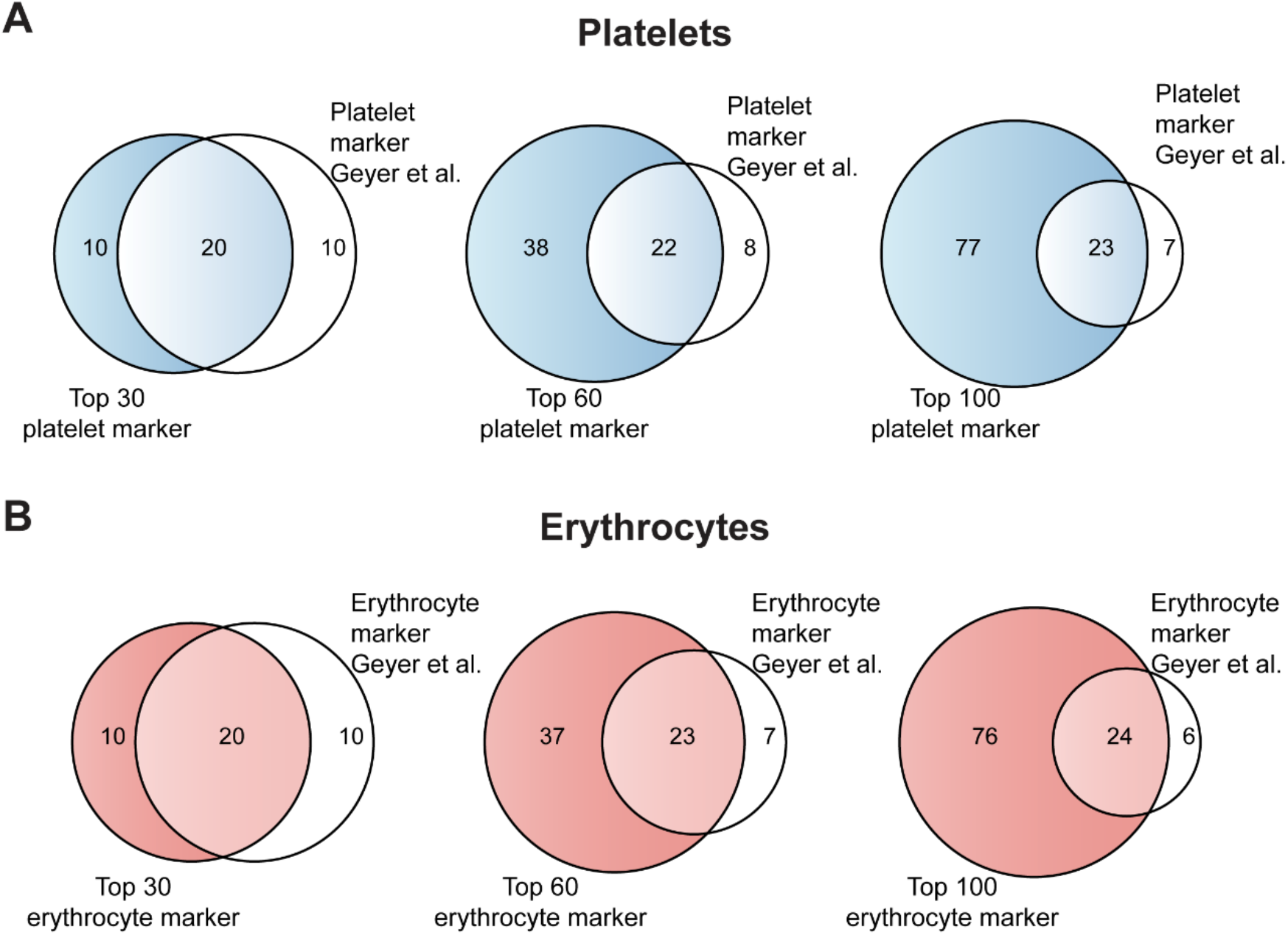
Validation and characterization of cell-specific quality markers. (A-B) Comparison of identified quality markers with previously published markers from Geyer et al. Venn diagrams show overlap between (A) platelet markers and (B) erythrocyte markers at three different stringency levels (top 30, top 60, and top 100 proteins). Despite differences in analytical platforms and experimental setup, substantial overlap is observed, with 67% identity (20 proteins) for both cell types in the top 30 panel, increasing to 22-23 proteins in the top 60 and 23-24 proteins in the top 100 markers.

**Supplementary Figure 2.**
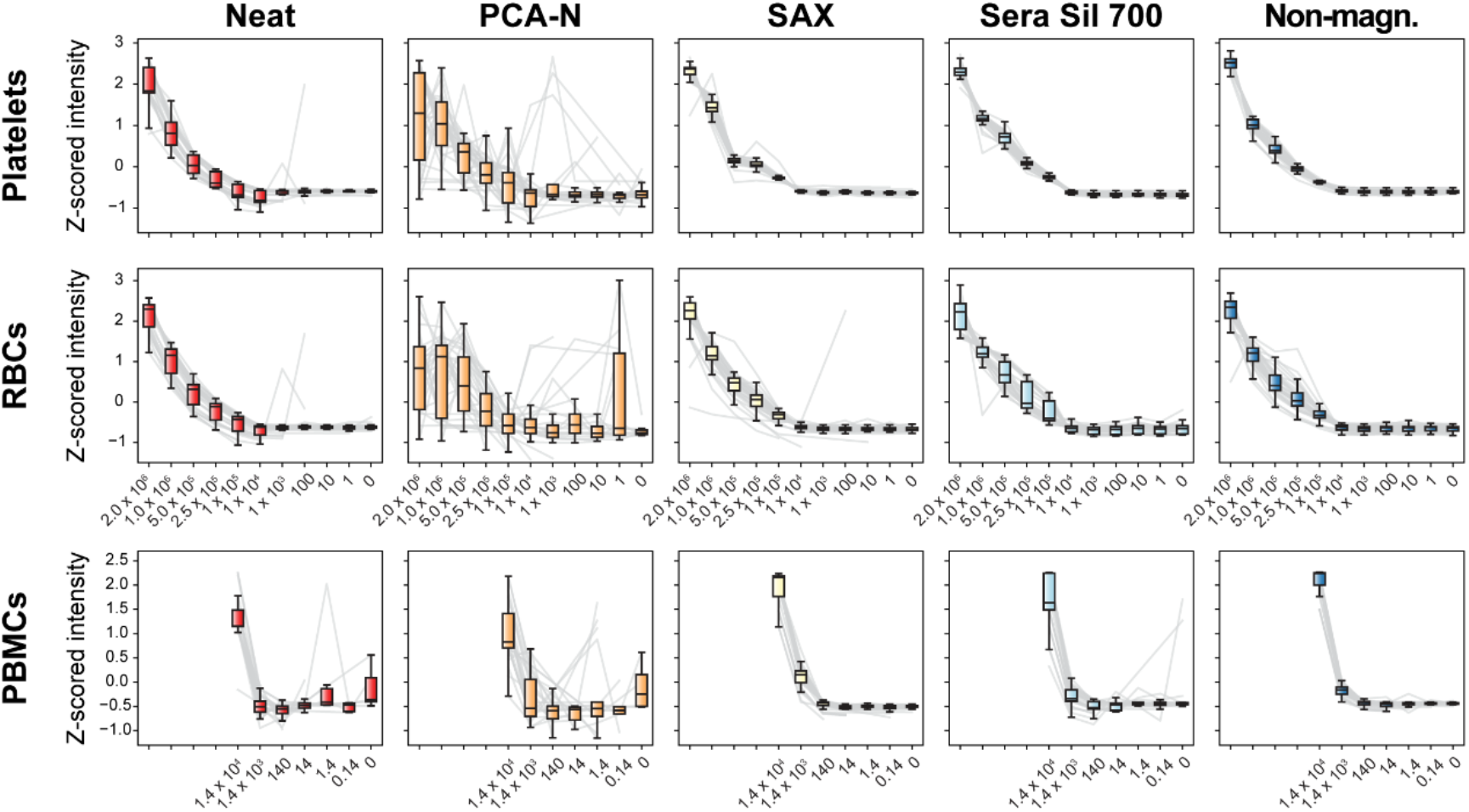
Z-scored intensity profiles of cell-specific quality markers across contamination series in five plasma proteomics workflows. Boxplots show Z-scored intensities of the top 30 quality markers for platelets (top row), erythrocytes (middle row), and PBMCs (bottom row) across all contamination levels and workflows. Gray lines represent individual marker proteins, while boxplots display the combined distribution of all 30 markers at each cell concentration. Data is Z-scored per workflow and per protein to allow direct comparison of concentration-dependent patterns.

**Supplementary Figure 3.**
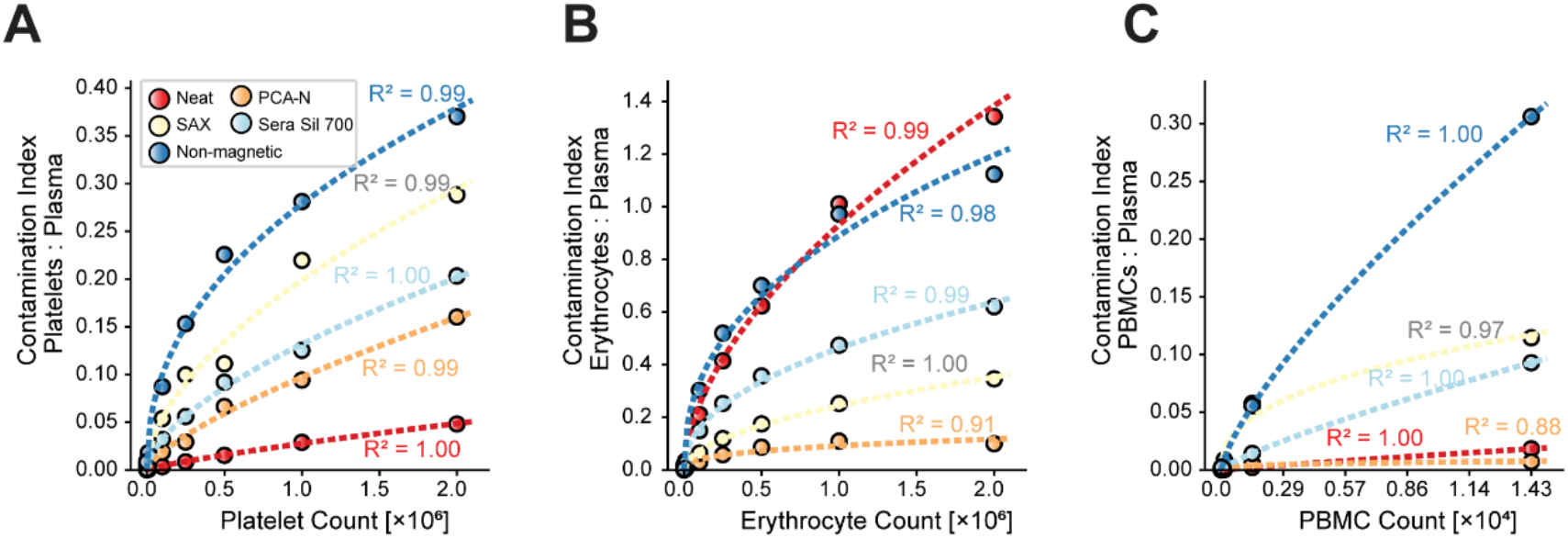
Power law model fitting of contamination index data across cell types. (A-C) Contamination index versus cell count with power law model fitting for (A) platelets, (B) erythrocytes, and (C) PBMCs across all five workflows. Points represent measured contamination indices at different cell counts, while curved lines show power law model fits. Coefficient of determination (R^2^) values are displayed for each workflow and cell type.

**Supplementary Figure 4.**
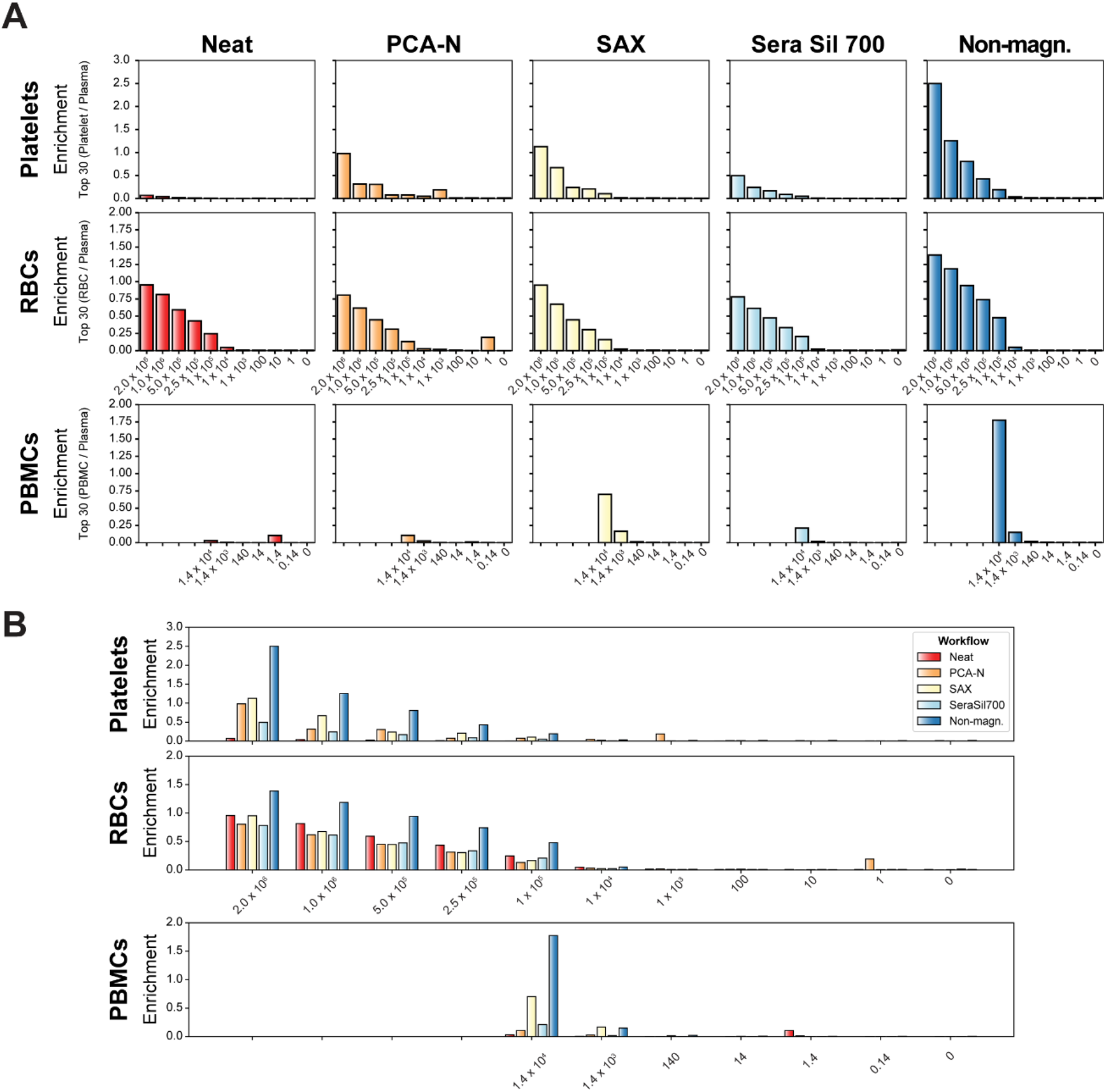
Enrichment scores of cell-specific markers across contamination series. (A) Enrichment scores for platelets (top row), erythrocytes (middle row), and PBMCs (bottom row) across all contamination levels for each of the five workflows. Enrichment score was calculated by dividing the summed intensity of the top 30 cell-specific markers by the summed intensity of the top 30 plasma proteins. (B) Direct comparison of enrichment scores across all five workflows for platelets (top), erythrocytes (middle), and PBMCs (bottom) at each contamination level.

**Supplementary Figure 5.**
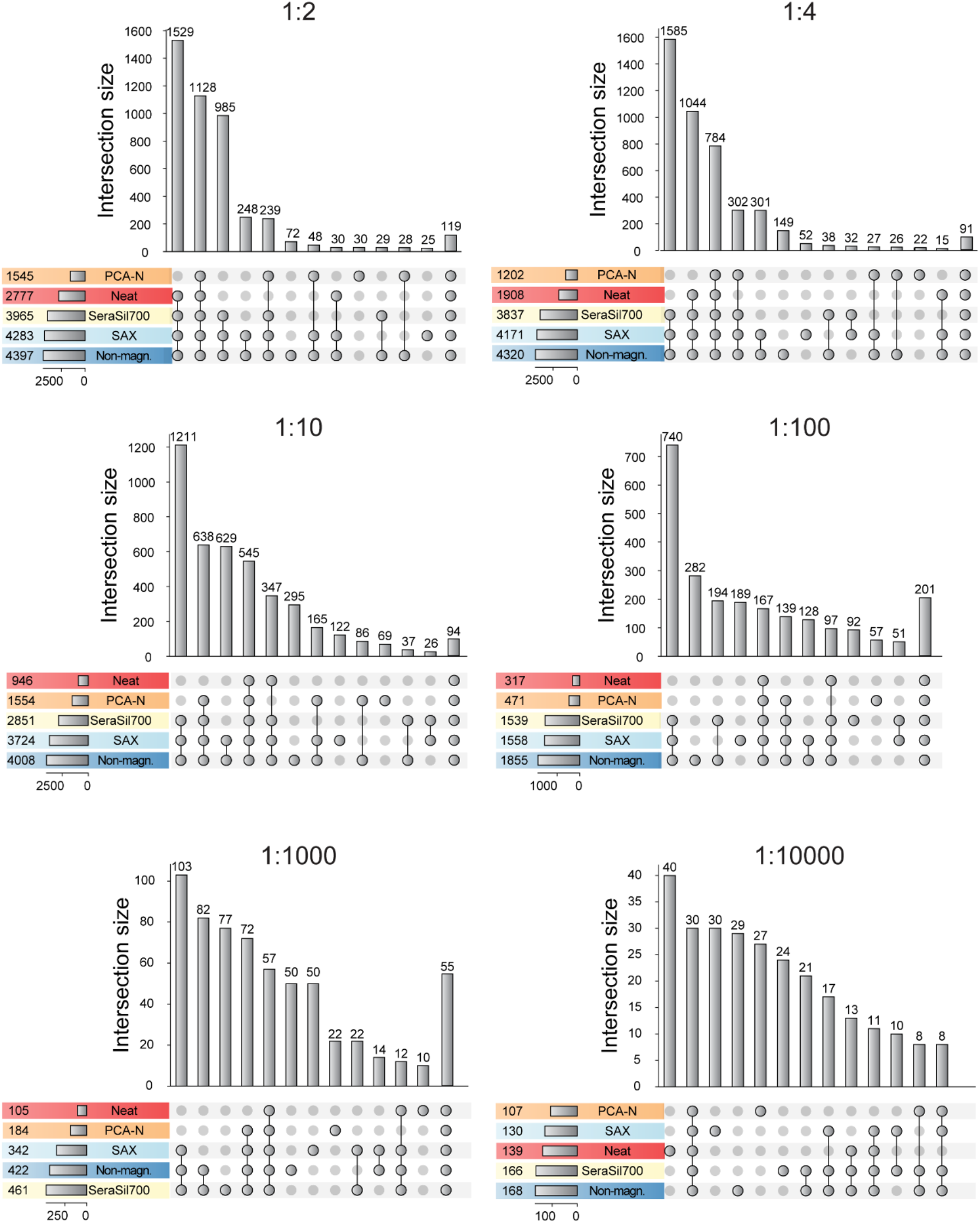
UpSet plots showing overlap of identified yeast proteins between workflows at different dilution ratios. Each panel represents a specific dilution ratio (1:2, 1:4, 1:10, 1:100, 1:1,000, and 1:10,000) with the total number of identified proteins per workflow shown in the horizontal bars on the left. The intersection size (vertical bars) indicates the number of proteins identified in each specific combination of workflows, with connected dots below showing which workflows contribute to each intersection. The total protein identifications decrease as the dilution ratio increases, with changing patterns of unique and shared identifications across workflows at different concentrations.

**Supplementary Figure 6.**
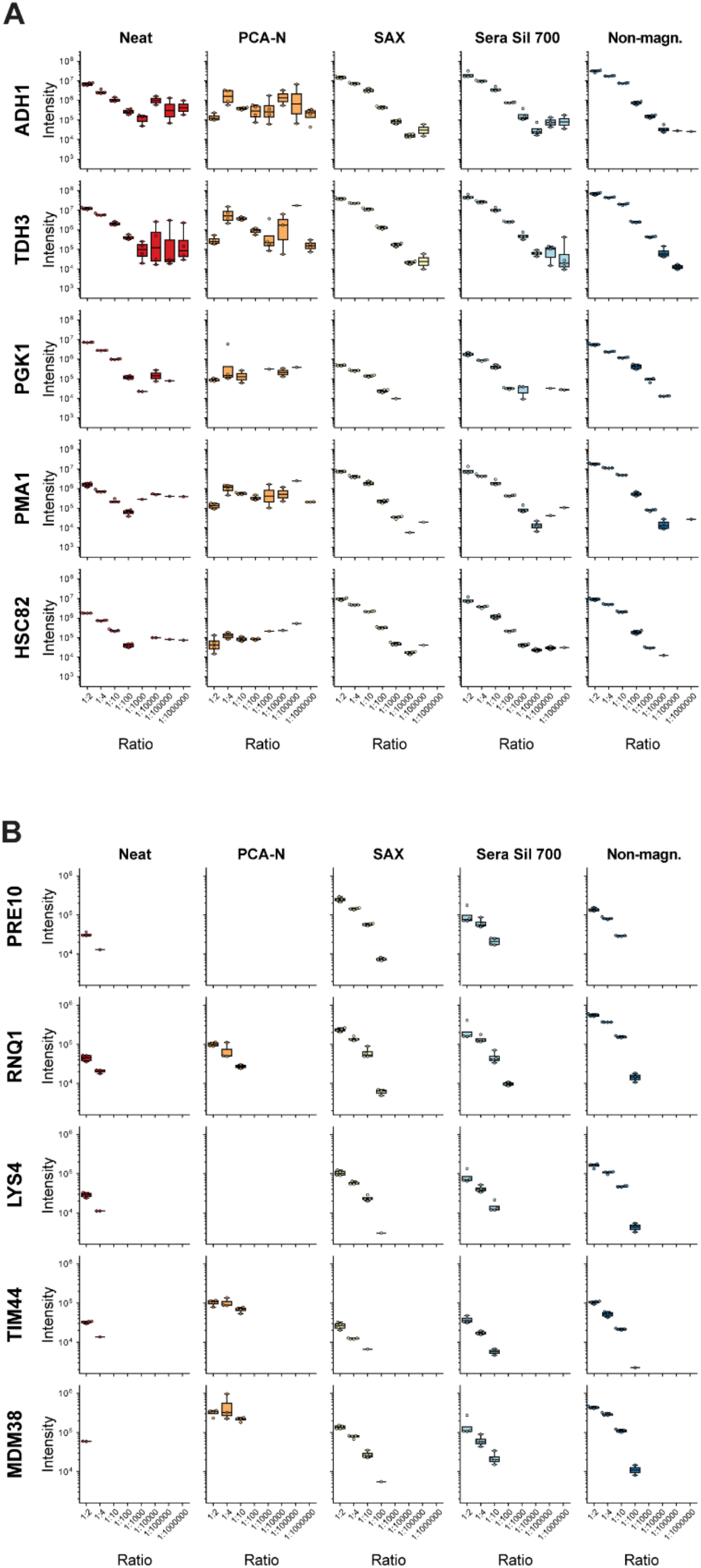
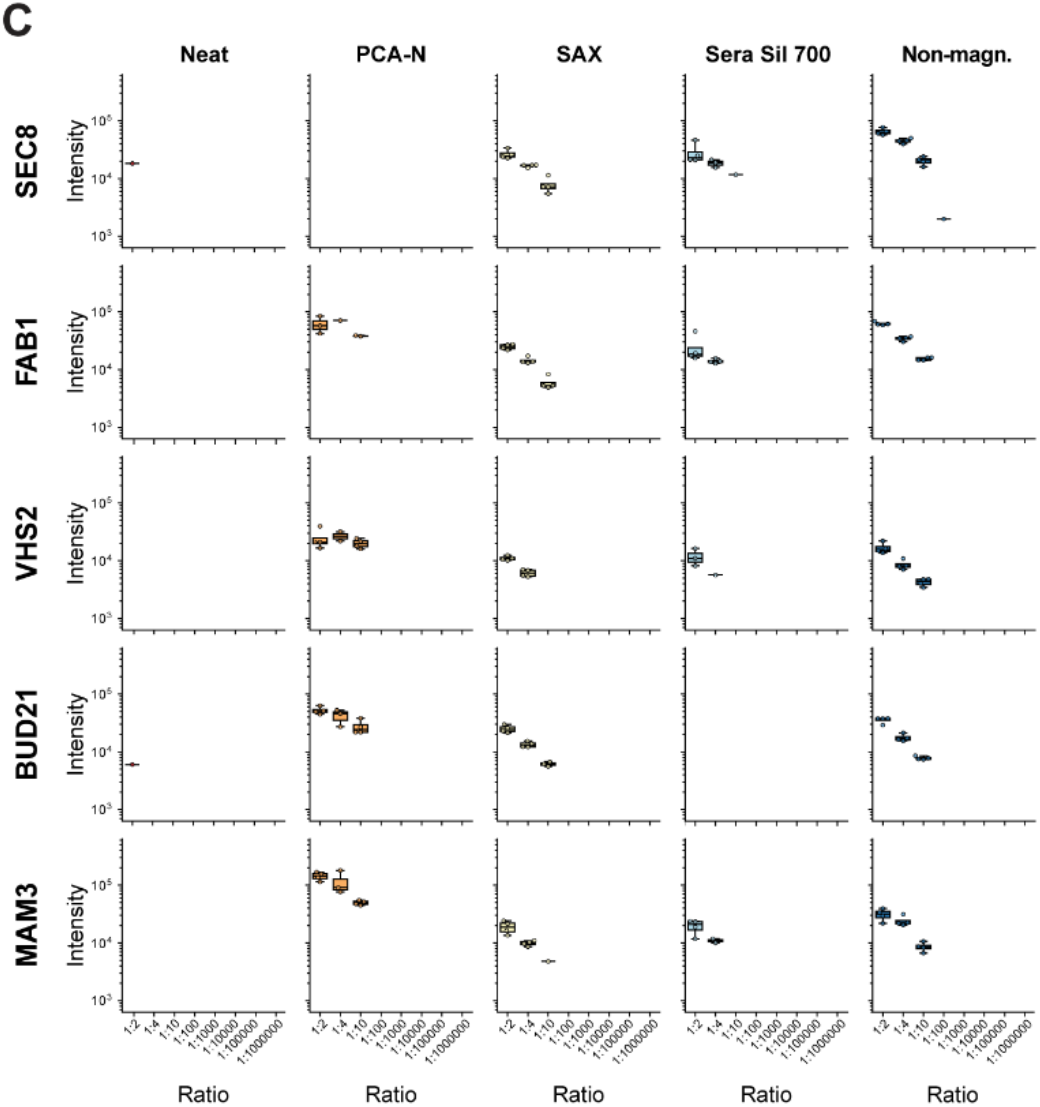
Dilution series analysis of yeast proteins across abundance ranges. (A) Representative high-abundant yeast proteins showing intensity measurements across dilution series from 1:2 to 1:10^6^ in all five workflows. (B) Representative medium-abundant yeast proteins displayed across the same dilution range and workflows. (C) Representative low-abundant yeast proteins shown across dilution series and workflows. Each panel displays protein intensity on the y-axis (log10 scale) against dilution ratio on the x-axis, with different colored lines representing each workflow’s detection profile for the specific proteins.

**Supplementary Figure 7.**
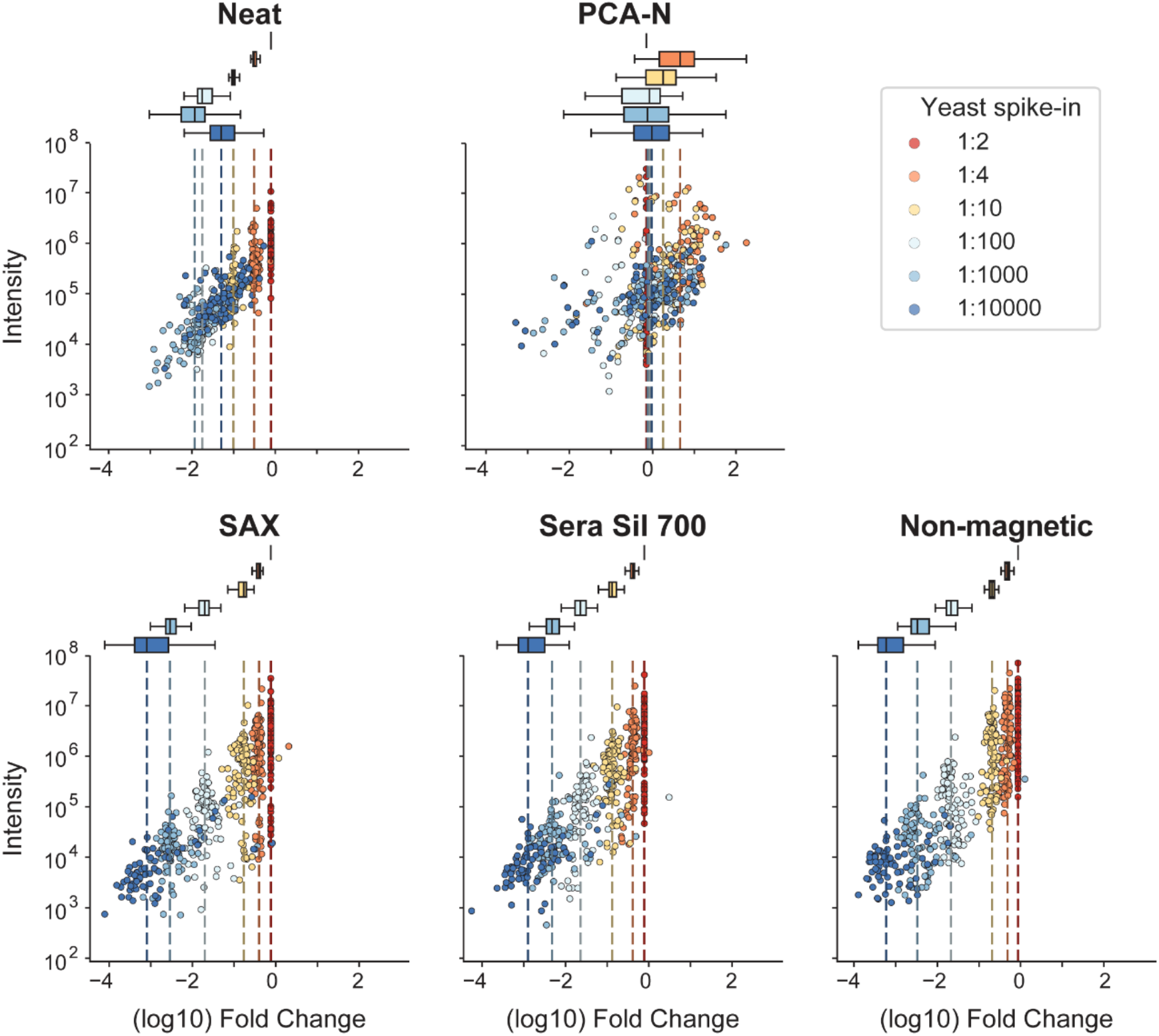
Analysis of protein enrichment across yeast protein abundance categories. Scatter plots displaying yeast protein intensity versus fold change relative to 1:2 (50%) spike-in for five plasma proteomics workflows: Neat, PCA-N, SAX, Sera Sil 700, and Non-magnetic. Data points are colored by dilution ratio. Boxplots at the top of each panel summarize fold change distributions at specific dilution points (marked by vertical dashed lines).

**Supplementary Figure 8.**
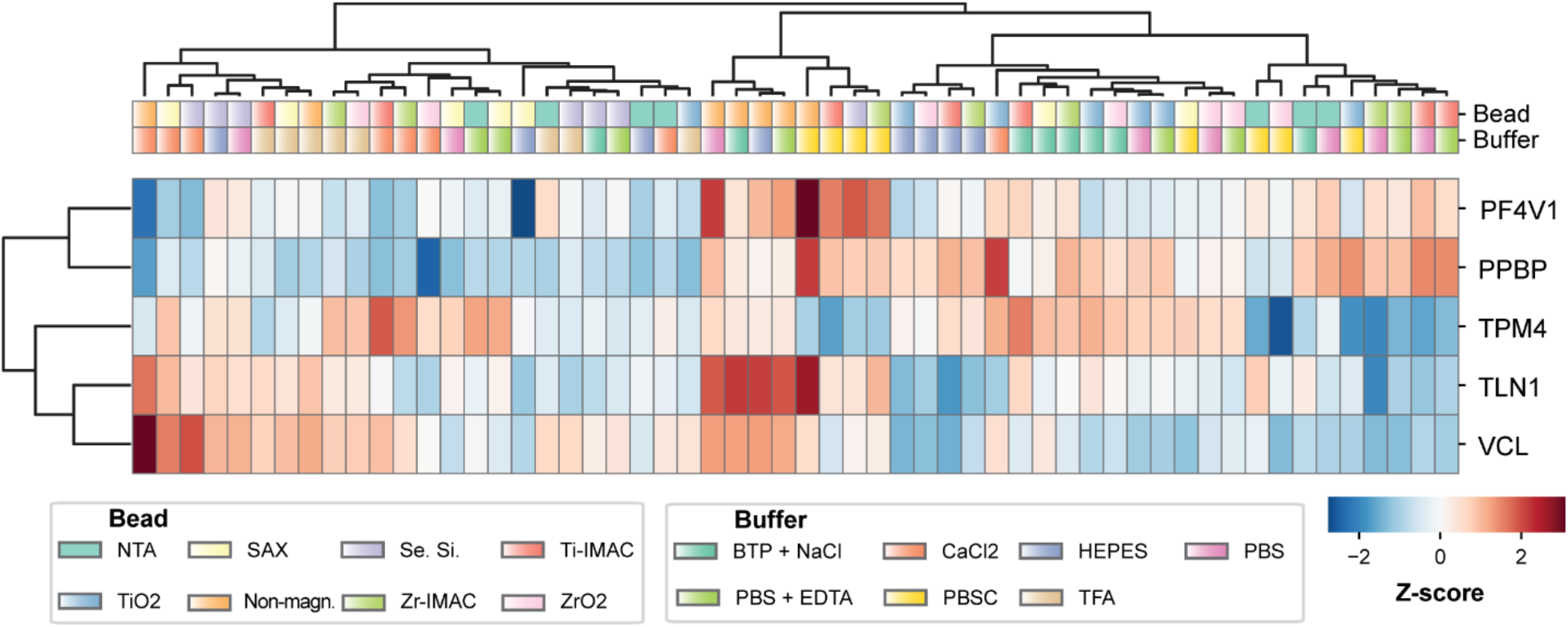
Analysis of platelet markers across bead-buffer combinations in platelet contaminated samples. Heatmap visualization of Z-scored intensities for five platelet marker proteins (PF4V1, PPBP, TPM4, TLN1, and VCL) across all tested bead-buffer combinations in platelet-contaminated plasma. Hierarchical clustering groups the markers based on similar behavior patterns across different experimental conditions. Bead types are indicated on the left and buffer conditions are shown at the bottom of the heatmap. Color intensity represents the relative abundance of each protein, with red indicating higher and blue indicating lower Z-scored intensities.

**Supplementary Figure 9.**
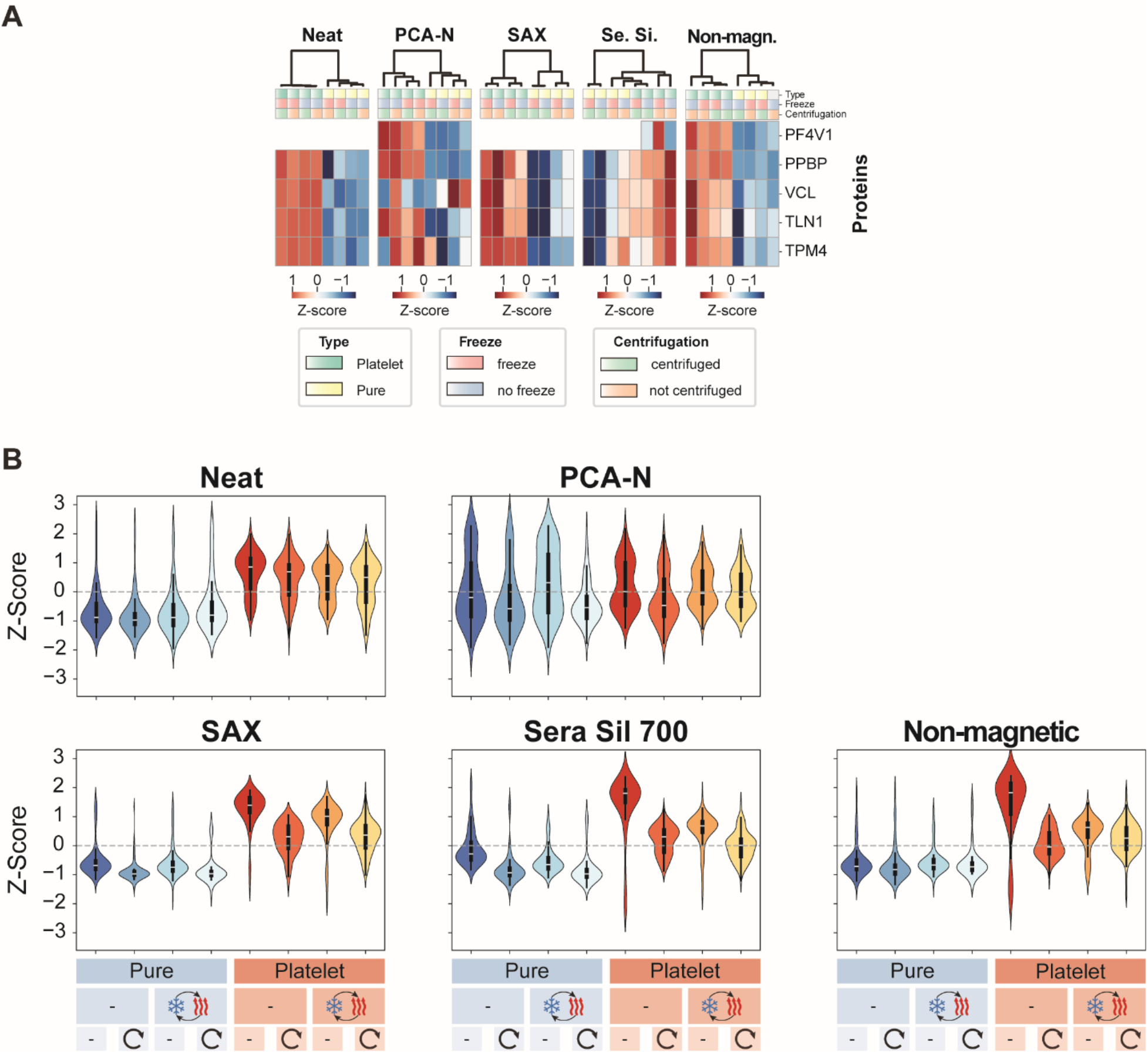
Platelet marker analysis across processing conditions. (A) Hierarchical clustering of five platelet marker proteins showing workflow-specific responses to processing steps. Data represents four replicates per condition. (B) Violin plots of Z-scored intensities for top 100 platelet markers across different workflows and processing conditions. Blue: pure plasma; Red: platelet-contaminated. Icons indicate sample type, freeze-thaw status, and centrifugation status.

**Supplementary Figure 10.**
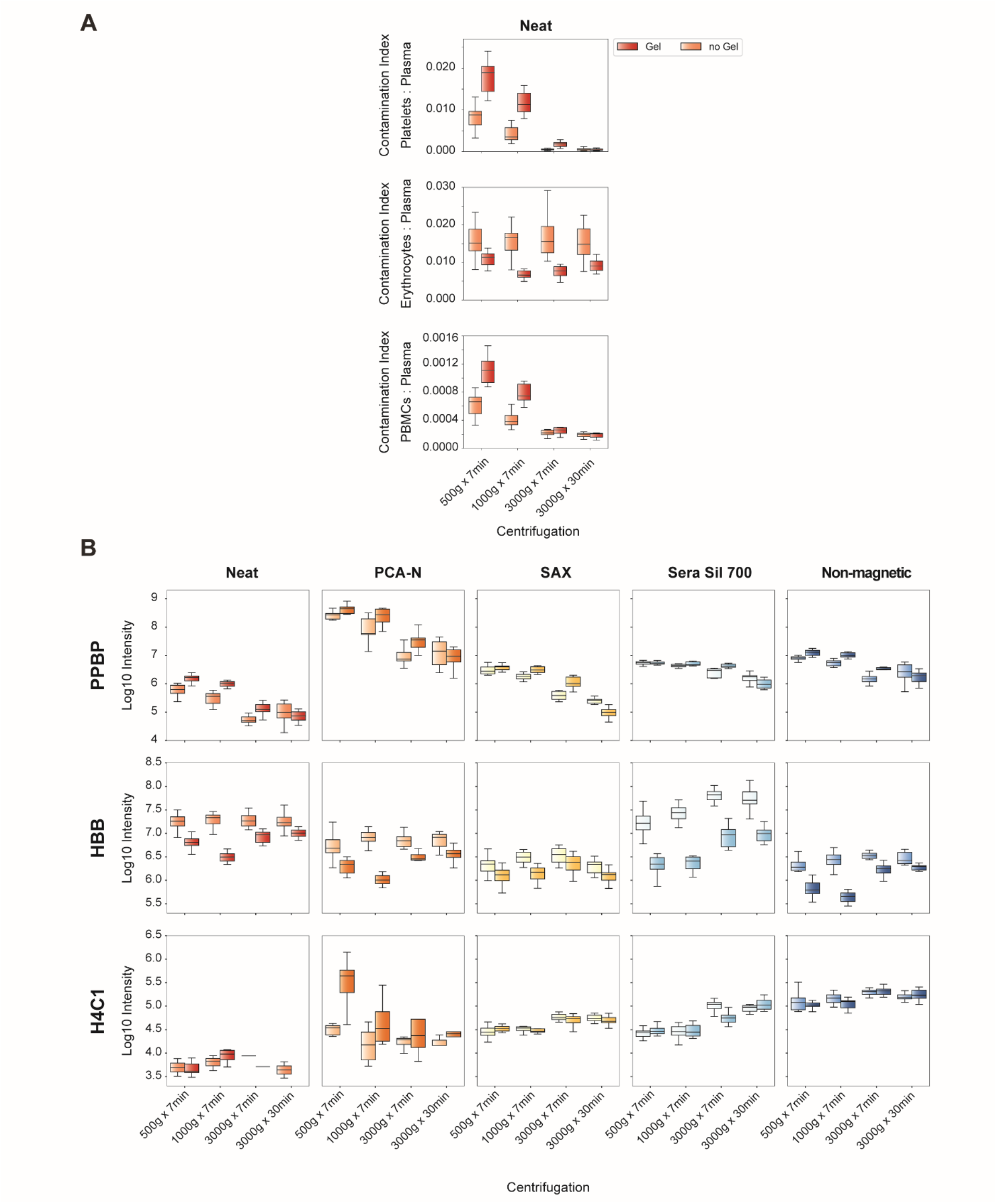
Analysis of contamination markers across centrifugation conditions. (A) Contamination indices for platelets, erythrocytes, and PBMCs in the neat workflow across all centrifugation conditions, comparing gel and no-gel tubes. (B) Abundance of specific marker proteins across centrifugation conditions and workflows. Top row: Platelet-specific marker PPBP (Platelet Basic Protein). Middle row: Erythrocyte-specific marker HBB (Hemoglobin Subunit Beta). Bottom row: PBMC-specific marker H4C1 (Histone H4). Boxplots show distribution of log10-transformed intensities across 11 individuals for each condition.

**Supplementary Figure 11.**
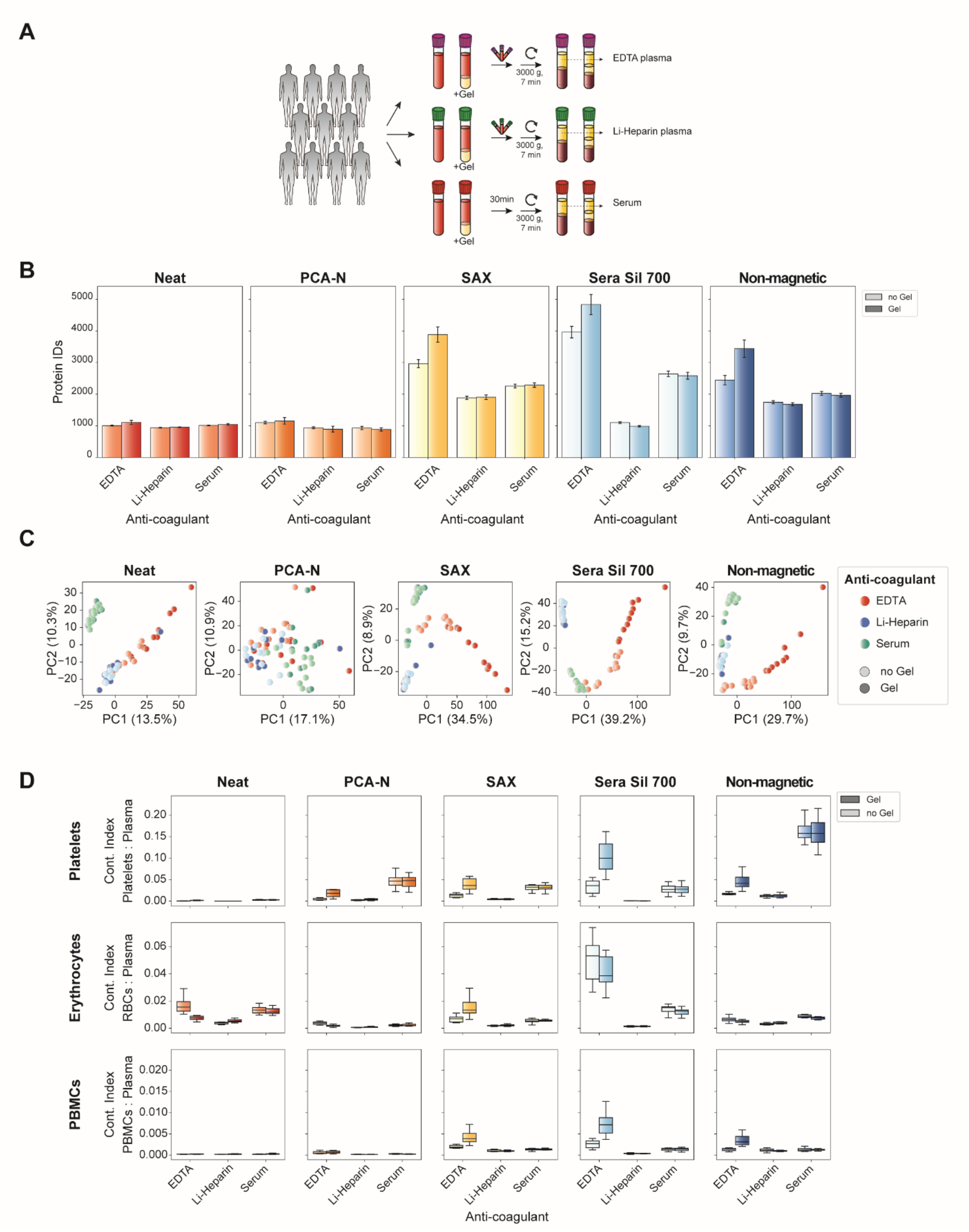
Impact of anti-coagulants on plasma proteome analysis. (A) Experimental design: Blood was collected from 11 healthy individuals into EDTA, Li-Heparin, and serum tubes, with and without gel separators. All samples were centrifuged at 3,000g for 7 min and processed through five proteomic workflows. (B) Number of protein identifications across five workflows for each anti-coagulant with and without gel separator tubes. (C) Principal component analysis of proteomics data for each workflow, colored by anti-coagulant type and tube presence. (D) Contamination indices for platelets, erythrocytes, and PBMCs across all workflows and anti-coagulant conditions.

## References

Aebersold R & Mann M (2016) Mass-spectrometric exploration of proteome structure and function. Nature 537: 347–355

Albrecht V, Müller-Reif JB, Brennsteiner V & Mann M (2025) A simplified perchloric acid workflow with neutralization (PCA-N) for democratizing deep plasma proteomics at population scale. 2025.03.24.645089 doi:10.1101/2025.03.24.645089 [PREPRINT]

Anderson NL & Anderson NG (2002) The human plasma proteome: history, character, and diagnostic prospects. Mol Cell Proteomics 1: 845–867

Anderson NL, Ptolemy AS & Rifai N (2013) The riddle of protein diagnostics: future bleak or bright? Clin Chem 59: 194–197

Bache N, Geyer PE, Bekker-Jensen DB, Hoerning O, Falkenby L, Treit PV, Doll S, Paron I, Müller JB, Meier F, et al (2018) A Novel LC System Embeds Analytes in Pre-formed Gradients for Rapid, Ultra-robust Proteomics *. Molecular & Cellular Proteomics 17: 2284–2296

Bader JM, Albrecht V & Mann M (2023) MS-Based Proteomics of Body Fluids: The End of the Beginning. Mol Cell Proteomics 22: 100577

Blume JE, Manning WC, Troiano G, Hornburg D, Figa M, Hesterberg L, Platt TL, Zhao X, Cuaresma RA, Everley PA, et al (2020) Rapid, deep and precise profiling of the plasma proteome with multi-nanoparticle protein corona. Nat Commun 11: 3662

Bowen RAR & Remaley AT (2014) Interferences from blood collection tube components on clinical chemistry assays. Biochem Med (Zagreb) 24: 31–44

Demichev V, Messner CB, Vernardis SI, Lilley KS & Ralser M (2020) DIA-NN: neural networks and interference correction enable deep proteome coverage in high throughput. Nat Methods 17: 41–44

Deutsch EW, Omenn GS, Sun Z, Maes M, Pernemalm M, Palaniappan KK, Letunica N, Vandenbrouck Y, Brun V, Tao S, et al (2021) Advances and Utility of the Human Plasma Proteome. J Proteome Res 20: 5241–5263

FDA-NIH Biomarker Working Group (2016) BEST (Biomarkers, EndpointS, and other Tools) Resource Silver Spring (MD): Food and Drug Administration (US)

Gao H, Zhan Y, Liu Y, Zhu Z, Zheng Y, Qian L, Xue Z, Cheng H, Nie Z, Ge W, et al (2025) Systematic evaluation of blood contamination in nanoparticle-based plasma proteomics. 2025.04.26.650757 doi:10.1101/2025.04.26.650757 [PREPRINT]

Geyer PE, Holdt LM, Teupser D & Mann M (2017) Revisiting biomarker discovery by plasma proteomics. Molecular Systems Biology 13: 942

Geyer PE, Kulak NA, Pichler G, Holdt LM, Teupser D & Mann M (2016) Plasma Proteome Profiling to Assess Human Health and Disease. Cell Syst 2: 185–195

Geyer PE, Voytik E, Treit PV, Doll S, Kleinhempel A, Niu L, Müller JB, Buchholtz M-L, Bader JM, Teupser D, et al (2019) Plasma Proteome Profiling to detect and avoid sample-related biases in biomarker studies. EMBO Mol Med 11: e10427

de Godoy LMF, Olsen JV, Cox J, Nielsen ML, Hubner NC, Fröhlich F, Walther TC & Mann M (2008) Comprehensive mass-spectrometry-based proteome quantification of haploid versus diploid yeast. Nature 455: 1251–1254

Guo T, Steen JA & Mann M (2025) Mass-spectrometry-based proteomics: from single cells to clinical applications. Nature 638: 901–911

Guzman UH, Martinez-Val A, Ye Z, Damoc E, Arrey TN, Pashkova A, Renuse S, Denisov E, Petzoldt J, Peterson AC, et al (2024) Ultra-fast label-free quantification and comprehensive proteome coverage with narrow-window data-independent acquisition. Nat Biotechnol 42: 1855–1866

Hendricks NG, Bhosale SD, Keoseyan AJ, Ortiz J, Stotland A, Seyedmohammad S, Nguyen CDL, Bui J, Moradian A, Mockus SM, et al (2024) An inflection point in high-throughput proteomics with Orbitrap Astral: analysis of biofluids, cells, and tissues. bioRxiv: 2024.04.26.591396

Hoofnagle AN, Whiteaker JR, Carr SA, Kuhn E, Liu T, Massoni SA, Thomas SN, Townsend RR, Zimmerman LJ, Boja E, et al (2016) Recommendations for the Generation, Quantification, Storage, and Handling of Peptides Used for Mass Spectrometry-Based Assays. Clin Chem 62: 48–69

Ignjatovic V, Geyer PE, Palaniappan KK, Chaaban JE, Omenn GS, Baker MS, Deutsch EW & Schwenk JM (2019) Mass Spectrometry-Based Plasma Proteomics: Considerations from Sample Collection to Achieving Translational Data. J Proteome Res 18: 4085–4097

Lancaster NM, Sinitcyn P, Forny P, Peters-Clarke TM, Fecher C, Smith AJ, Shishkova E, Arrey TN, Pashkova A, Robinson ML, et al (2024) Fast and deep phosphoproteome analysis with the Orbitrap Astral mass spectrometer. Nat Commun 15: 7016

Lee JY, Kim JY, Park GW, Cheon MH, Kwon K-H, Ahn YH, Moon MH, Lee H-J, Paik YK & Yoo JS (2011) Targeted mass spectrometric approach for biomarker discovery and validation with nonglycosylated tryptic peptides from N-linked glycoproteins in human plasma. Mol Cell Proteomics 10: M111.009290

Li J, Zhao C, Liu Y, Zhang W & Qin W (2024) An Ultra-Deep Quantitative Plasma Proteomics Strategy. doi:10.26434/chemrxiv-2024-lcmhl [PREPRINT]

Messner CB, Demichev V, Wendisch D, Michalick L, White M, Freiwald A, Textoris-Taube K, Vernardis SI, Egger A-S, Kreidl M, et al (2020) Ultra-High-Throughput Clinical Proteomics Reveals Classifiers of COVID-19 Infection. Cell Syst 11: 11-24.e4

Mischak H, Allmaier G, Apweiler R, Attwood T, Baumann M, Benigni A, Bennett SE, Bischoff R, Bongcam-Rudloff E, Capasso G, et al (2010) Recommendations for biomarker identification and qualification in clinical proteomics. Sci Transl Med 2: 46ps42

Niu L, Stinson SE, Holm LA, Lund MAV, Fonvig CE, Cobuccio L, Meisner J, Juel HB, Fadista J, Thiele M, et al (2025) Plasma proteome variation and its genetic determinants in children and adolescents. Nat Genet 57: 635–646

Rai AJ, Gelfand CA, Haywood BC, Warunek DJ, Yi J, Schuchard MD, Mehigh RJ, Cockrill SL, Scott GBI, Tammen H, et al (2005) HUPO Plasma Proteome Project specimen collection and handling: towards the standardization of parameters for plasma proteome samples. Proteomics 5: 3262–3277

Robinson WH, Lindstrom TM, Cheung RK & Sokolove J (2013) Mechanistic biomarkers for clinical decision making in rheumatic diseases. Nat Rev Rheumatol 9: 267–276

Sadgrove NJ, Batra JK & Batra S (2024) Critical Insights on Preparation of Platelet-Rich Plasma in Tubes With a Thixotropic Gel Separator. J Drugs Dermatol 23: 979–985

Schrohl A-S, Wuürtz S, Kohn E, Banks RE, Nielsen HJ, Sweep FCGJ & Bruünner N (2008) Banking of Biological Fluids for Studies of Disease-associated Protein Biomarkers. Molecular & Cellular Proteomics 7: 2061–2066

Serrano LR, Peters-Clarke TM, Arrey TN, Damoc E, Robinson ML, Lancaster NM, Shishkova E, Moss C, Pashkova A, Sinitcyn P, et al (2024) The One Hour Human Proteome. Mol Cell Proteomics 23: 100760

Singh S, Dodt J, Volkers P, Hethershaw E, Philippou H, Ivaskevicius V, Imhof D, Oldenburg J & Biswas A (2019) Structure functional insights into calcium binding during the activation of coagulation factor XIII A. Sci Rep 9: 11324

Skates SJ, Gillette MA, LaBaer J, Carr SA, Anderson L, Liebler DC, Ransohoff D, Rifai N, Kondratovich M, Težak Ž, et al (2013) Statistical design for biospecimen cohort size in proteomics-based biomarker discovery and verification studies. J Proteome Res 12: 5383–5394

Stewart HI, Grinfeld D, Giannakopulos A, Petzoldt J, Shanley T, Garland M, Denisov E, Peterson AC, Damoc E, Zeller M, et al (2023) Parallelized Acquisition of Orbitrap and Astral Analyzers Enables High-Throughput Quantitative Analysis. Anal Chem 95: 15656–15664

Surinova S, Schiess R, Hüttenhain R, Cerciello F, Wollscheid B & Aebersold R (2011) On the development of plasma protein biomarkers. J Proteome Res 10: 5–16

Timms JF, Arslan-Low E, Gentry-Maharaj A, Luo Z, T’Jampens D, Podust VN, Ford J, Fung ET, Gammerman A, Jacobs I, et al (2007) Preanalytic influence of sample handling on SELDI-TOF serum protein profiles. Clin Chem 53: 645–656

Viode A, Smolen KK, van Zalm P, Stevenson D, Jha M, Parker K, IMPACC Network‡, Levy O, Steen JA & Steen H (2024) Longitudinal plasma proteomic analysis of 1117 hospitalized patients with COVID-19 identifies features associated with severity and outcomes. Sci Adv 10: eadl5762

Viode A, van Zalm P, Smolen KK, Fatou B, Stevenson D, Jha M, Levy O, Steen J, Steen H, & ON BEHALF OF THE IMPACC NETWORK (2023) A simple, time- and cost-effective, high-throughput depletion strategy for deep plasma proteomics. Science Advances 9: eadf9717

Wu CC, Tsantilas KA, Park J, Plubell D, Sanders JA, Naicker P, Govender I, Buthelezi S, Stoychev S, Jordaan J, et al (2024) Mag-Net: Rapid enrichment of membrane-bound particles enables high coverage quantitative analysis of the plasma proteome. bioRxiv: 2023.06.10.544439

